# Radioproteomics stratifies molecular response to antifibrotic treatment in pulmonary fibrosis

**DOI:** 10.1101/2024.03.27.586923

**Authors:** David Lauer, Cheryl Yael Magnin, Luca Kolly, Huijuan Wang, Matthias Brunner, Mamta Charbria, Grazia Maria Cereghetti, Hubert Gabryś, Stephanie Tanadini-Lang, Anne-Christine Uldry, Manfred Heller, Stijn E Verleden, Kerstin Klein, Adela-Cristina Sarbu, Manuela Funke-Chambour, Lukas Ebner, Oliver Distler, Britta Maurer, Janine Gote-Schniering

## Abstract

Antifibrotic therapy with nintedanib is the clinical mainstay in the treatment of progressive fibrosing interstitial lung disease (ILD). High-dimensional medical image analysis, known as radiomics, provides quantitative insights into organ-scale pathophysiology, generating digital disease fingerprints. Here, we used an integrative analysis of radiomic and proteomic profiles (radioproteomics) to assess whether changes in radiomic signatures can stratify the degree of antifibrotic response to nintedanib in (experimental) fibrosing ILD. Unsupervised clustering of delta radiomic profiles revealed two distinct imaging phenotypes in mice treated with nintedanib, contrary to conventional densitometry readouts, which showed a more uniform response. Integrative analysis of delta radiomics and proteomics demonstrated that these phenotypes reflected different treatment response states, as further evidenced on transcriptional and cellular levels. Importantly, radioproteomics signatures paralleled disease- and drug related biological pathway activity with high specificity, including extracellular matrix (ECM) remodeling, cell cycle activity, wound healing, and metabolic activity. Evaluation of the preclinical molecular response-defining features, particularly those linked to ECM remodeling, in a cohort of nintedanib-treated fibrosing ILD patients, accurately stratified patients based on their extent of lung function decline. In conclusion, delta radiomics has great potential to serve as a non-invasive and readily accessible surrogate of molecular response phenotypes in fibrosing ILD. This could pave the way for personalized treatment strategies and improved patient outcomes.

## Introduction

Fibrotic remodeling of the lung interstitium is the shared pathomechanism across various interstitial lung diseases (ILDs) of different etiologies, including idiopathic pulmonary fibrosis (IPF) and connective tissue disease (CTD)-associated ILD as the most prevalent subtypes. For patients with a progressive fibrosing (PF-ILD) phenotype, treatment with the antifibrotic multitarget tyrosine kinase inhibitor nintedanib is recommended ^1^. While nintedanib has proven effective in slowing pulmonary function decline in multiple clinical trials, it comes with a relatively high rate of side effects ^2–4^. Consequently, there is a pressing need to assess treatment efficacy and identify individuals who may not benefit from therapy early in the disease course.

Current evaluation of treatment response primarily relies on longitudinal lung function measurements, which are prone to intra-subject variability, can be influenced by extrapulmonary parameters, and lack insights into the underlying molecular response ^5,6^. Liquid- or tissue-derived readouts could partially address these limitations, but validated biomarkers are not yet available and repeated lung biopsies are not a viable option due to the associated interventional risks ^7^. Radiomics analysis of routinely performed high-resolution computed tomography (HRCT) scans has great potential to serve as a non-invasive solution for evaluation of treatment response in individual patients in four dimensions (3D space + time) ^8,9^. Radiomic features are computationally retrieved, quantitative data extracted from radiological imaging data, which describe the tissue in terms of its intensity, texture and shape properties ^10^, thus creating digital disease fingerprints ^11^. The added value compared to conventional visual radiological analysis or quantitative characterization methodologies such as CALIPER ^12^, lies in their ability to capture image phenotypes beyond human visual perception^13^, thereby aiming to close the gap between patient screening and precision medicine ^14^.

Radiomics is based on the premise that the underlying pathophysiology is reflected in the imaging phenotype and that radiomic features can quantify these links, offering insights into organ-scale pathophysiology. Previous work including our own has shown that radiomics can convey morphologic and molecular tissue characteristics with important implications for personalized diagnosis and prognostication ^15–18^. Delta radiomics, which quantifies the feature variation between two imaging time points, and thus captures longitudinal phenotypic changes, has emerged as a method to predict and quantify treatment response in various types of cancer ^19–21^. Its potential for the stratification of antifibrotic treatment response in (progressive) fibrosing ILD has not yet been studied.

This study aimed to evaluate whether delta radiomics can be used to stratify the degree of molecular response to antifibrotic treatment with nintedanib using the well-established bleomycin-induced lung fibrosis model.

Unsupervised clustering of delta radiomic profiles revealed two distinct imaging phenotypes in mice treated with nintedanib, despite conventional CT-derived lung densitometric readouts suggesting a uniform response. Radioproteomics demonstrated that these phenotypes reflected different treatment response states, which we could confirm by immunofluorescence and gene expression analysis. Importantly, we discovered that radioproteomic association modules paralleled distinct disease-related biological pathway activities and cell type signatures. Evaluation of the preclinical response-defining features in a nintedanib-treated PF-ILD cohort accurately stratified patients according to their extent of lung function decline. Collectively, our analyses demonstrate the ability of delta radiomics to non-invasively stratify molecular response to anti-fibrotic treatment in experimental fibrosing ILD and indicate its potential for application in human PF-ILD.

## Methods

A detailed description of the methods is provided in the supplementary material.

### Animal experimentation and ethics statement

Lung-derived delta radiomic profiles were studied upon treatment with nintedanib in the well-established murine bleomycin-induced fibrosing ILD model ^22,23^. Briefly, lung fibrosis was induced in C57BL/6J mice (*n*=30, female, 8-weeks old) by intratracheal instillation with 2 U/kg bleomycin sulfate^17^. Mice were randomized into study groups and treatment with 60 mg/kg nintedanib (*n*=15) or vehicle-only (deionized water, *n*=15) was provided once daily *per os* (p.o.) from day 7-20 in a double-blinded manner. Lung microCT scans (SkyScan 1176; Bruker, Kontich, Belgium) of each animal were acquired pre-(day 7) and post-treatment (day 21) as previously described ^17^. All mice were sacrificed 24 hours after the final treatment, followed by exsanguination, transcardial perfusion, and collection of the lung tissue for molecular analysis. Approval for animal experimentation was granted by the cantonal veterinary office (license number ZH082/2021) and experimentation was performed in strict compliance with Swiss animal protection laws. Mice were excluded from further analysis if humane endpoints were reached (*n*=3) or if severe lung abnormalities, including atelectasis or unilateral fibrosis development, were evident on microCT scans (*n*=3). The final sample size for nintedanib- and vehicle-treated mice was n=10 and n=14, respectively.

### Patient cohort, clinical data, and ethics statement

We validated our experimental findings in a retrospectively selected PF-ILD cohort of 19 patients from the Bern University Hospital, Bern, Switzerland and the SWISS-IIP cohort that were undergoing treatment with nintedanib. Approval for the study was granted by the local ethics committee (BASEC-ID: 2023-01920 [ILDALMO]; PB_2016_01524 [SWISS-IIP]). A total of 359 patients were screened for the following eligibility criteria: a) diagnosis of progressive fibrosing ILD ^24^, including IPF, systemic-sclerosis associated ILD (SSc-ILD), hypersensitivity pneumonitis (HP), or drug-induced ILD, b) treatment with nintedanib (≥100 mg twice daily; ≥6 months), c) availability of pre- and post-treatment HRCT scans fulfilling the predefined quality criteria (supplementary methods), d) pre- and post-treatment pulmonary function test (PFT) recording fulfilling the predefined quality criteria (supplementary material), e) absence of secondary lung diseases at times of HRCT and PFT recordings. In total, 54 out 359 patients received nintedanib treatment for ≥6 months, with 19 fulfilling also the remaining inclusion criteria. Summaries of patient demographics and clinical characteristics, and the HRCT scan acquisition parameters are provided in **Table 1** and **Supplementary Table 1**, respectively.

**Table 1.**
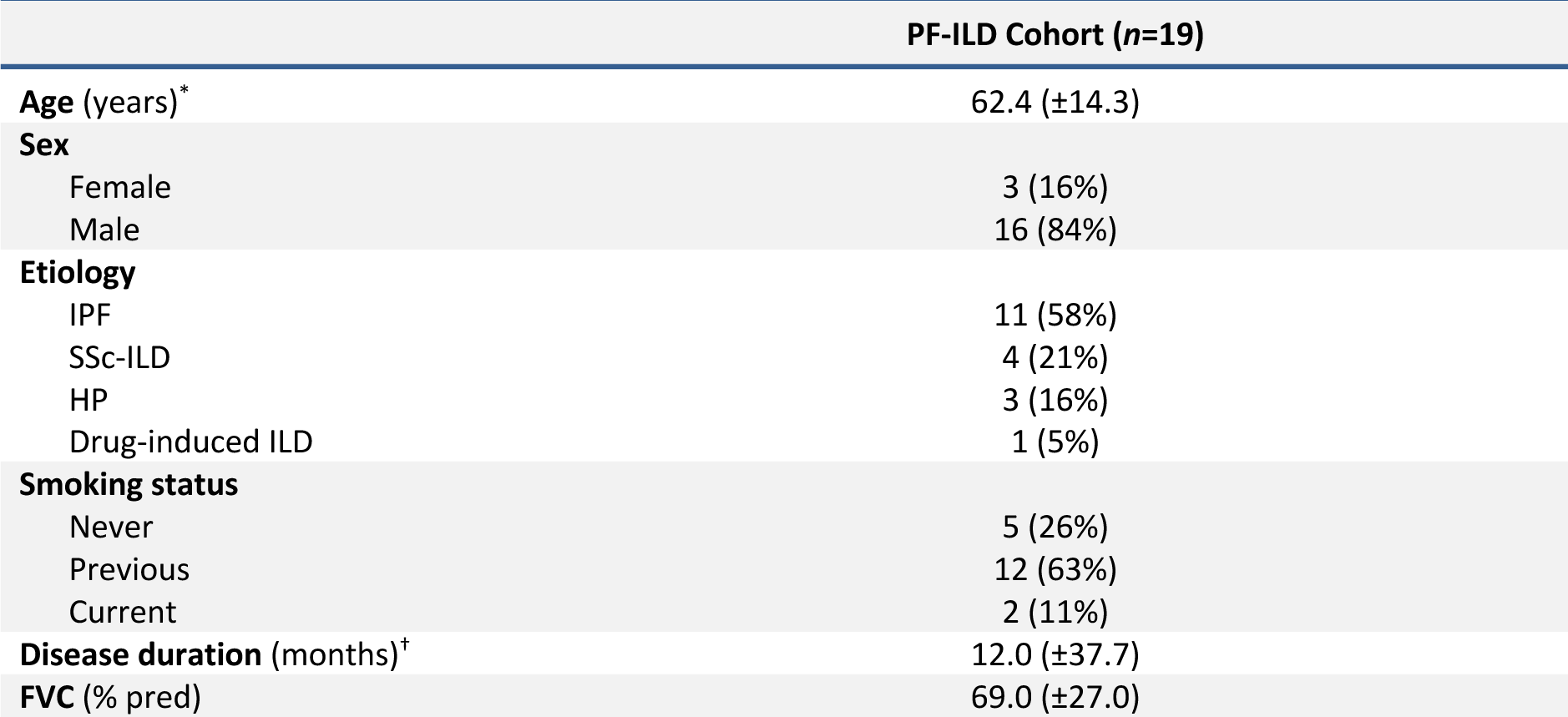
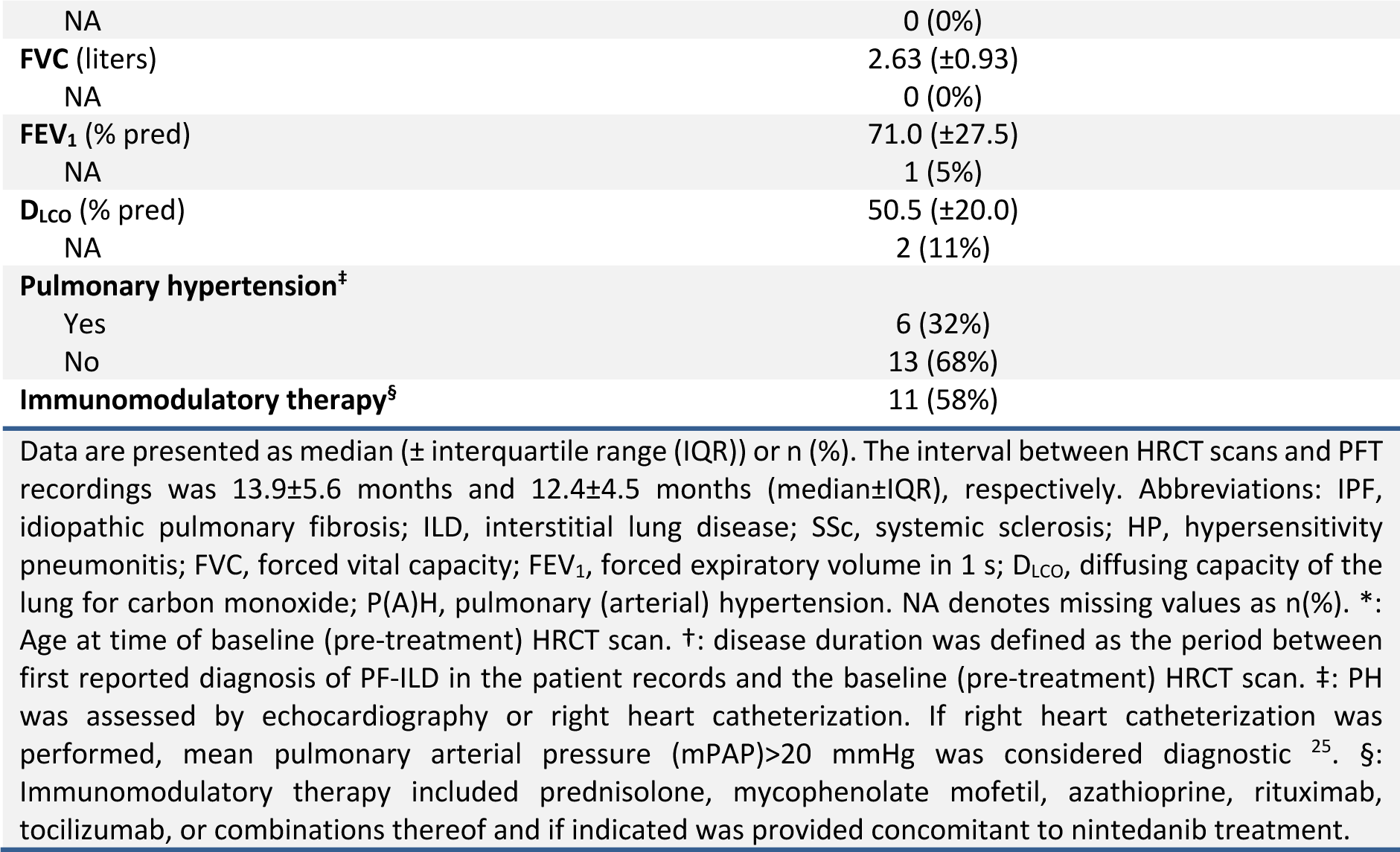
Summary of the clinical parameters of the PF-ILD cohort at baseline.

| PF-ILD Cohort ($n=19$) | |
| --- | --- |
| **Age (years)\*** | 62.4 ($\pm 14.3$) |
| **Sex** |  |
| Female | 3 (16%) |
| Male | 16 (84%) |
| **Etiology** |  |
| IPF | 11 (58%) |
| SSc-ILD | 4 (21%) |
| HP | 3 (16%) |
| Drug-induced ILD | 1 (5%) |
| **Smoking status** |  |
| Never | 5 (26%) |
| Previous | 12 (63%) |
| Current | 2 (11%) |
| **Disease duration (months)<sup>†</sup>** | 12.0 ($\pm 37.7$) |
| **FVC (% pred)** | 69.0 ($\pm 27.0$) |
| NA | 0 (0%) |
| **FVC (liters)** | 2.63 (±0.93) |
| NA | 0 (0%) |
| **FEV<sub>1</sub> (% pred)** | 71.0 (±27.5) |
| NA | 1 (5%) |
| **D<sub>LCO</sub> (% pred)** | 50.5 (±20.0) |
| NA | 2 (11%) |
| **Pulmonary hypertension<sup>‡</sup>** |  |
| Yes | 6 (32%) |
| No | 13 (68%) |
| **Immunomodulatory therapy<sup>§</sup>** | 11 (58%) |

Data are presented as median (± interquartile range (IQR)) or n (%). The interval between HRCT scans and PFT recordings was 13.9±5.6 months and 12.4±4.5 months (median±IQR), respectively. Abbreviations: IPF, idiopathic pulmonary fibrosis; ILD, interstitial lung disease; SSs, systemic sclerosis; HP, hypersensitivity pneumonitis; FVC, forced vital capacity; FEV<sub>1</sub>, forced expiratory volume in 1 s; D<sub>LCO</sub>, diffusing capacity of the lung for carbon monoxide; P(A)H, pulmonary (arterial) hypertension. NA denotes missing values as n(%). \*: Age at time of baseline (pre-treatment) HRCT scan. †: disease duration was defined as the period between first reported diagnosis of PF-ILD in the patient records and the baseline (pre-treatment) HRCT scan. ‡: PH was assessed by echocardiography or right heart catheterization. If right heart catheterization was performed, mean pulmonary arterial pressure (mPAP)>20 mmHg was considered diagnostic <sup>25</sup>. §: Immunomodulatory therapy included prednisolone, mycophenolate mofetil, azathioprine, rituximab, tocilizumab, or combinations thereof and if indicated was provided concomitant to nintedanib treatment.

### Delta Radiomics Calculation

Calculation of radiomic features was performed on semi-automatically segmented lungs using Z-Rad software (v.7.3.1, https://medical-physics-usz.github.io/) as previously described ^17^. Mouse lungs were resized to isotropic voxels of 0.15 mm. To achieve a comparable voxel size in patients, human lungs were resized to isotropic voxels of 2.75 mm, corresponding to an estimated 6000-fold volumetric difference ^26^. Both mouse and human lung volumes were discretized to a fixed bin size of 50 HU in a range of -1000 HU to 200 HU. From the resized volumes, 1’388 radiomic features were calculated per lung scan and time point, corresponding to histogram (*n*=17), texture (*n*=137), shape (*n*=2), and wavelet-transformed features (*n*=1’232). Delta radiomic features describing the change of each feature between pre-and post-treatment, were expressed as delta values: ΔFeature = Feature (t_2_) – Feature (t_1_) ^27^. Lung density measurement was inferred from the radiomic lung attenuation histogram-derived feature *hist_mean*.

### Proteomics and Phosphoproteomics

For proteomics and phosphoproteomics, the middle lobe of the right mouse lung was snap frozen in liquid nitrogen and stored at -80°C until processing. Sample preparation and mass spectrometry profiling was performed at the Proteomics & Mass Spectrometry Core Facility (PMSCF) at the University of Bern using standard protocols. For comparative proteomics, all vehicle-(*n*=14) and nintedanib-treated (*n*=10) samples were analyzed. One vehicle sample was excluded from analysis due to issues in sample preparation. Differential expression of proteins between groups of interest was calculated in R using the “limma” package with standard settings. For phosphoproteomics, randomly selected subsets of vehicle-(*n*=5) and nintedanib-treated (*n*=5) were analyzed. Differential expression of phosphosites and subsequent kinase activity enrichment analysis (KAEA) was performed as described in ^28^.

### Gene expression analysis

Total RNA was isolated from blood-free cranial lobes of the right mouse lung using the RNeasy Tissue Mini Kit (Qiagen, Hombrechtikon, Switzerland). Isolated RNA was reverse-transcribed into cDNA using the Transcriptor First Strand cDNA Synthesis Kit (Roche Diagnostics, Switzerland). Expression of selected Nintedanib target genes was analyzed by SYBR Green quantitative PCR as described in ^29^. Expression of mRNA was expressed to delta Ct values with *Rplp0* as reference gene. Fold changes were calculated using the delta-delta Ct method. A list of the primer pairs is provided in **Supplementary Table 2**.

### Immunofluorescence and microscopy

Formalin-fixed paraffin-embedded lung sections at 3 µm thickness were deparaffinized, followed by heat-mediated antigen retrieval and blocking for nonspecific antibody binding with 5% BSA. Incubation with primary antibodies was performed overnight at 4°C, followed by incubation with secondary fluorescence-labeled antibodies for 2 h at room temperature. Nuclei were visualized by counterstaining with 4’,6-diamidino-2-phenylindole (DAPI) for 10 min at RT. Antibodies and dilutions are listed in **Supplementary Table 3**. Microscopic imaging was performed with an AxioScan.Z1 slide scanner (Zeiss, Feldbach, Switzerland) using a Plan-Apochromat 20x/0.8 M27 objective. Cells positively stained for α-SMA were quantified using the “Positive cell detection” tool of the open source software QuPath (v.0.4.0). From each sample, five representative areas at 500×500 µm were analyzed and the sample average was used for statistical analyses.

### Unsupervised clustering

Unsupervised agglomerative hierarchical or k-means clustering of z-scored features was performed to identify subgroups of mice or patients with similar delta radiomic feature patterns. Clusterability was evaluated by Hopkin’s statistic H. The optimal number of clusters was determined by average silhouette statistics. Stability of clusters was assessed by Jaccard bootstrapping.

### Variable importance evaluation

The importance of each delta radiomic feature cluster assignment by unsupervised clustering was calculated by filter-based variable importance, retaining only features with classification score≥0.9 (**Supplementary Table 4**).

### Radioproteomic Correlation Analysis

Spearman’s rank correlation coefficient ⍴ was calculated between delta radiomic features subsets and the log2-transformed expression intensity of every protein, retaining only proteins with p<0.05 and ⍴≥0.6 to establish radioproteomic association modules. Pearson’s correlation coefficient r was calculated between delta radiomic features subsets and the fraction of α-SMA positive cells.

### Gene Ontology and Reactome Pathway Enrichment

Lists of differentially expressed (DE) proteins or radiomics-correlated proteins were entered into Gene Ontology (GO) or Reactome pathway enrichment analysis, retaining results after false discovery rate adjustment (p<0.05).

### Cell Type Signature Enrichment Analysis

To infer relative cell type frequency changes between two groups from proteomics data, we applied signature enrichment analysis as described in ^30,31^, utilizing their single cell marker gene dataset. Cell type signatures were defined as sets of genes with cell-type specific gene expression of log2 fold change>0.3 and adjusted p<0.05 (**Supplementary Table 5**).

### Statistical Analyses

All statistical analyses were performed in R (v.4.3.1.) environment. For all analyses, a p<0.05 was considered statistically significant unless stated otherwise.

## Results

### Delta radiomics uncovers heterogeneity in antifibrotic drug response

To study the effects of antifibrotic treatment on radiomic signatures, we collected microCT-derived radiomic features in mice with bleomycin-induced lung fibrosis (*n*=24) before (day 7) and after (day 21) treatment with nintedanib (*n*=10) or vehicle (*n*=14) (**Figure 1A**). The change in feature expression between pre- and post-treatment was quantified as delta radiomics. We considered only variables that were stable (ICC≥0.75) against semi-automatic lung segmentation and excluded highly correlated (Spearman’s ⍴≥0.85) features, resulting in a final set of 244 delta radiomic features that entered analysis (**Supplementary Figure 1A-C**).

**Figure 1.**
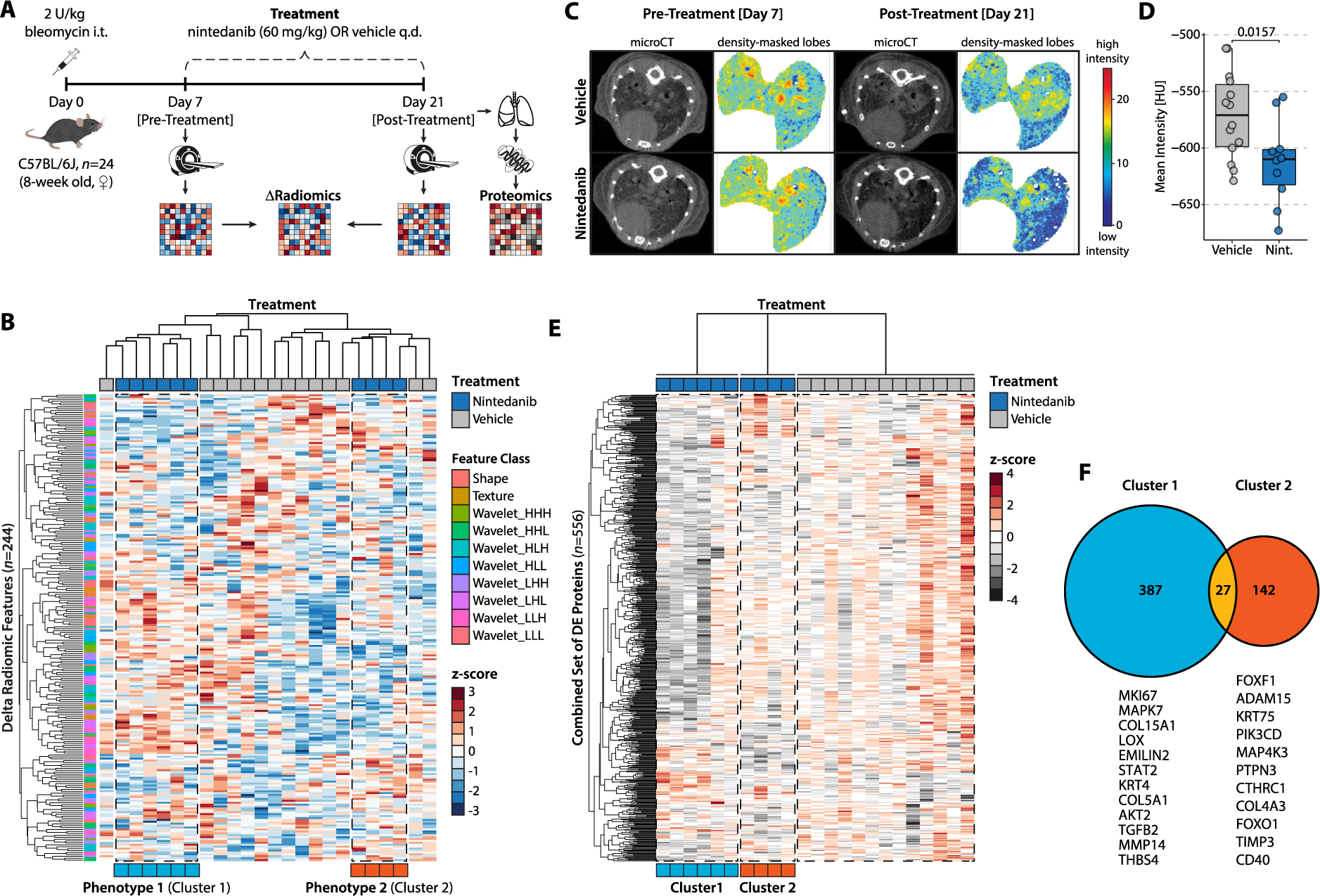
Delta radiomics uncovers heterogeneity in antifibrotic drug response. (**A**) Experiment schematic. C57BL/6J mice with bleomycin-induced lung fibrosis received treatment with nintedanib (*n*=10) or vehicle (*n*=14). Lung microCT scans were acquired of each mouse pre- (day 7) and post-treatment (day 21) for analysis of radiomic measures. The change in radiomic feature expression was expressed as delta radiomics. Lung tissue was collected 24 hours after the final treatment application for molecular analyses. (**B**) Heatmap displaying the results of unsupervised hierarchical clustering of z-scored delta radiomic features (*n*=244) in all mice. Treatment groups and the class of each delta radiomic variable are indicated. (**C**) Representative lung microCT images and matching density-masked lobes of nintedanib- and vehicle-treated mice with bleomycin-induced lung fibrosis at pre- and post-treatment level. (**D**) Lung tissue density expressed as mean Hounsfield unit (HU) intensity post-treatment. Unpaired Student’s t-test was used to compare the groups. (**E**) Heatmap showing the expression profiles of the combined set of DE proteins (*n*=556) in cluster 1 and cluster 2 compared to vehicle-treated mice. Log2-transformed protein expression values were z-scored. (**F**) Venn diagram depicting the number of differentially expressed proteins in cluster 1 (*n*=414) and cluster 2 (*n*=169) compared to the vehicle group. Selected DE proteins unique to cluster 1 or cluster 2 with functions implicated in disease pathophysiology are denoted.

Unsupervised hierarchical clustering of delta radiomics revealed heterogeneous response profiles in nintedanib-treated mice, highlighting the presence of two distinct imaging phenotypes (n_cluster1_=6, n_cluster2_=4) (**Figure 1B**). Subanalysis by k-means clustering confirmed their statistical stability (Jaccard coefficients>0.90) (**Supplementary Figures 1D-E**). Intriguingly, these clusters were not discernible through conventional lung densitometry, which showed a significantly (p=0.0157, unpaired Student’s t test) reduced tissue density in response to nintedanib treatment, consistent with previous reports ^23,32^ (**Figures 1C-D, Supplementary Figure 1F**). Untargeted phosphoproteome quantification in a subset of vehicle and nintedanib-treated mice 24 hours after the final treatment further confirmed successful and homogeneous target engagement with suppression of key drug-related pathways based on kinase activity enrichment analysis, including MTOR and MAP2K1 signaling ^33,34^ (**Supplementary Figures 1G-H**), thus affirming the efficiency of the drug treatment.

To evaluate if the two delta radiomic clusters differ on molecular level, we performed proteomics analysis. Differential expression analysis of the 7006 identified proteins in cluster 1 and 2 against the vehicle group uncovered substantial differences between the two delta radiomics phenotypes. While 414 proteins (373 downregulated and 41 upregulated) were differentially expressed in cluster 1, only 169 proteins (127 downregulated and 42 upregulated) showed differential expression in cluster 2 compared to vehicle-treated mice (**Supplementary Figures 1I-J, Supplementary Tables 6-7**). Most notably, only minor differentially expressed (DE) protein overlap (5%) was observed between the two clusters (**Figures 1E-F**), suggesting that different molecular response phenotypes are captured by delta radiomics.

### Delta radiomic phenotypes reflect differences in molecular response to antifibrotic treatment

To describe the underlying biology of the two delta radiomic clusters in closer detail, we analyzed the differences on a molecular and cellular level. On protein level, 386 proteins were differentially regulated between the two clusters (**Figure 2A, Supplementary Table 8**). Gene Ontology (GO) mapping of the downregulated proteins (*n*=269) revealed enrichment of terms related to pro-fibrotic activity, including extracellular matrix (ECM) organization, regulation of cell growth, and fibroblast proliferation (**Figure 2B**, **Supplementary Table 9**). In contrast, the upregulated proteins (*n*=117) were enriched for pathways related to wound healing and tissue regeneration, including epithelial cell migration and hemostasis (**Figure 2B**, **Supplementary Table 10**).

**Figure 2.**
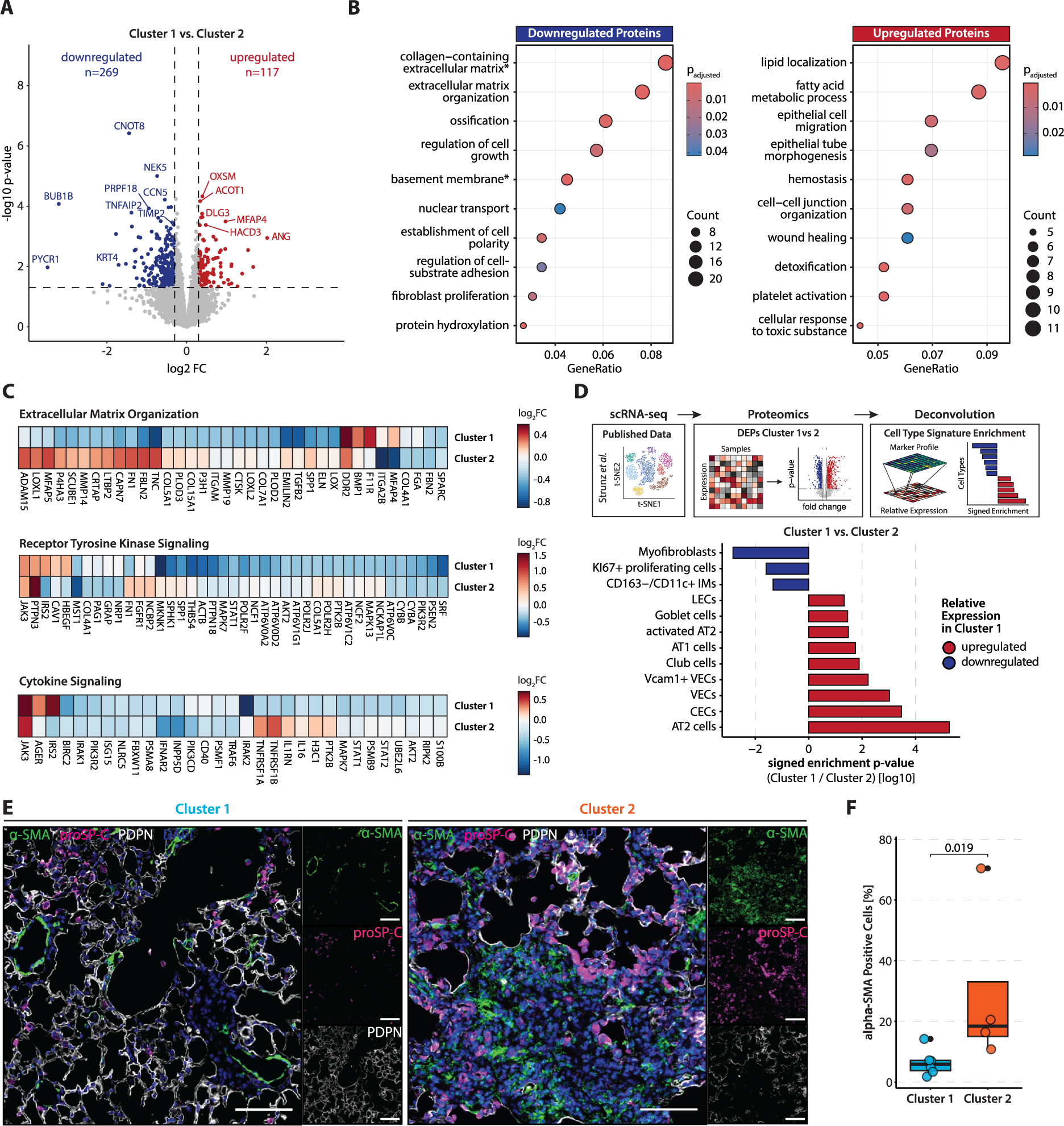
Delta radiomic phenotypes reflect molecular response to antifibrotic treatment. (**A**) Volcano plot of the DE proteins between clusters 1 and 2. Proteins with log2FC>0.3 and p<0.05 were considered significantly different. Down- and upregulated proteins are highlighted in blue and red, respectively. (**B**) GO pathway analysis of the down- and upregulated DE proteins. Terms marked with an asterisk are of cellular compartment (CC) ontology, all others are of biological process (BP) ontology. (**C**) Heatmap of DE proteins included in Reactome pathways “extracellular matrix organization”, “receptor tyrosine kinase signaling”, and “cytokine signaling”, and their expression in clusters 1 and 2 compared to vehicle-treated mice. (**D**) Analysis workflow and bar chart depicting the results from cell type deconvolution analysis. The change of the indicated cell type signature between clusters 1 and 2 is expressed as signed log10 enrichment p-value. (**E**) Representative immunofluorescence stainings of fibrotic regions in clusters 1 and 2. Images show nuclei (DAPI), AT2 cells (proSP-C), myofibroblasts (α-SMA), and AT1 cells (PDPN). Regions are 500×500 µm in size, scale bar 100 µm. (**F**) Percentage of α-SMA+ and proSP-C+ cells in fibrotic regions of cluster 1 and cluster 2 samples. Mann-Whitney U-test was used to compare the groups.

Furthermore, targeted activity analysis of pathways known to be modulated by nintedanib, including extracellular matrix (ECM) organization, receptor tyrosine kinase (RTK) signaling, and cytokine signaling ^35^, revealed more extensive pathway inhibition in cluster 1 (**Figure 2C**). Whereas most targets involved in ECM organization (e.g. COL5A1, COL12A1, TNC) and remodeling (e.g. MMP2, MMP14, TIMP2, LOX) were downregulated in cluster 1 compared to vehicle, their expression was less changed in cluster 2. Similarly, proteins involved in RTK (e.g. SPP1, STAT1, AKT2, MAPK7, MAPK13) and cytokine signaling (IL6, IRAK1, IRAK2, and PIK3R2) showed higher suppression in cluster 1 than cluster 2. To independently validate our proteomics results, we performed quantitative PCR of selected gene targets of nintedanib. Aligning with the proteomic observations, we found significant (p<0.05, unpaired Student’s t-test) suppression of pro-fibrotic (*Col1a1*, *Col3a1*, *Fn1*), pro-inflammatory (*Il6*, *Spp1*), and nintedanib-targeted (*Tgfb1*, *Cxcl1*, *Tnf*, *Cd40l*) transcripts in cluster 1 compared to cluster 2 (**Supplementary Figure 2**).

Preclinical studies demonstrated that nintedanib inhibits myofibroblast differentiation ^22^, cell proliferation ^23^, and macrophage activation ^36^, thereby promoting regeneration of alveolar epithelial cells. To interrogate the cluster-specific effects of nintedanib on the cellular level, we performed cell type deconvolution analysis of our proteomics data as described in ^30^. This technique quantifies the enrichment of single cell RNA-sequencing (scRNA-seq)-derived cell type marker signatures in bulk cell analysis data such as proteomics or transcriptomics, allowing to estimate cell type frequency changes between two conditions. Deconvolution revealed lower levels of myofibroblasts, interstitial macrophages, and KI-67+ proliferating cells along with a higher fraction of alveolar type II (AT2) and type I (AT1) lung epithelial cells in cluster 1 compared to cluster 2 (**Figure 2D**). Tissue immunofluorescence staining for the myofibroblast marker α-SMA together with the AT1 marker Podoplanin (PDPN) and the AT2 marker proSP-C confirmed a significant (p=0.019, Mann-Whitney U test) lower abundance of α-SMA+ myofibroblast infiltrates in fibrotic regions in samples of cluster 1 compared to cluster 2 (**Figure 2E-F**).

Taken together, we found that delta radiomics-defined treatment sub-clusters exhibited distinct molecular and cellular characteristics, suggesting a higher degree of response to nintedanib in cluster 1.

### Delta radiomic features reflect changes in disease-relevant molecular pathway activity

Having established that delta radiomic phenotypes are able to characterize the extent of molecular response to nintedanib treatment, we next investigated the contribution of individual features to non-invasively convey pathway-specific molecular information. To do so, we first identified features promoting cluster separation by analysis of univariate variable importance, resulting in 54 variables with a classification score≥0.90 (**Figure 3A**, **Supplementary Figure 3A-C**). For each of these features, we then established radioproteomic association modules (*n*=54) by determining the respective correlating protein sets (Spearman’s |ρ|≥0.6, p<0.05) in a sample-matched, cluster-independent approach. Pathway annotation of positively or negatively correlated proteins revealed significant enrichment of Reactome terms for 45 features, covering 367 unique pathways (**Figure 3B, Supplementary Table 11**). These findings were replicated through GO:BP database annotation (**Supplementary Figure 3D, Supplementary Table 12**). Importantly, subsets of association modules were highly distinctive towards specific disease pathophysiology-related pathway activity, including ECM remodeling, cell cycle activity, wound healing, or metabolic processes. K-means sub-clustering of nintedanib-treated samples on features positively correlating with ECM organization (*n*=8) or hemostasis (*n*=7) reproduced the original two clusters, thereby indicating suppression of ECM remodeling as well as promotion of wound healing in cluster 1 (**Supplementary Figure 3E**).

**Figure 3.**
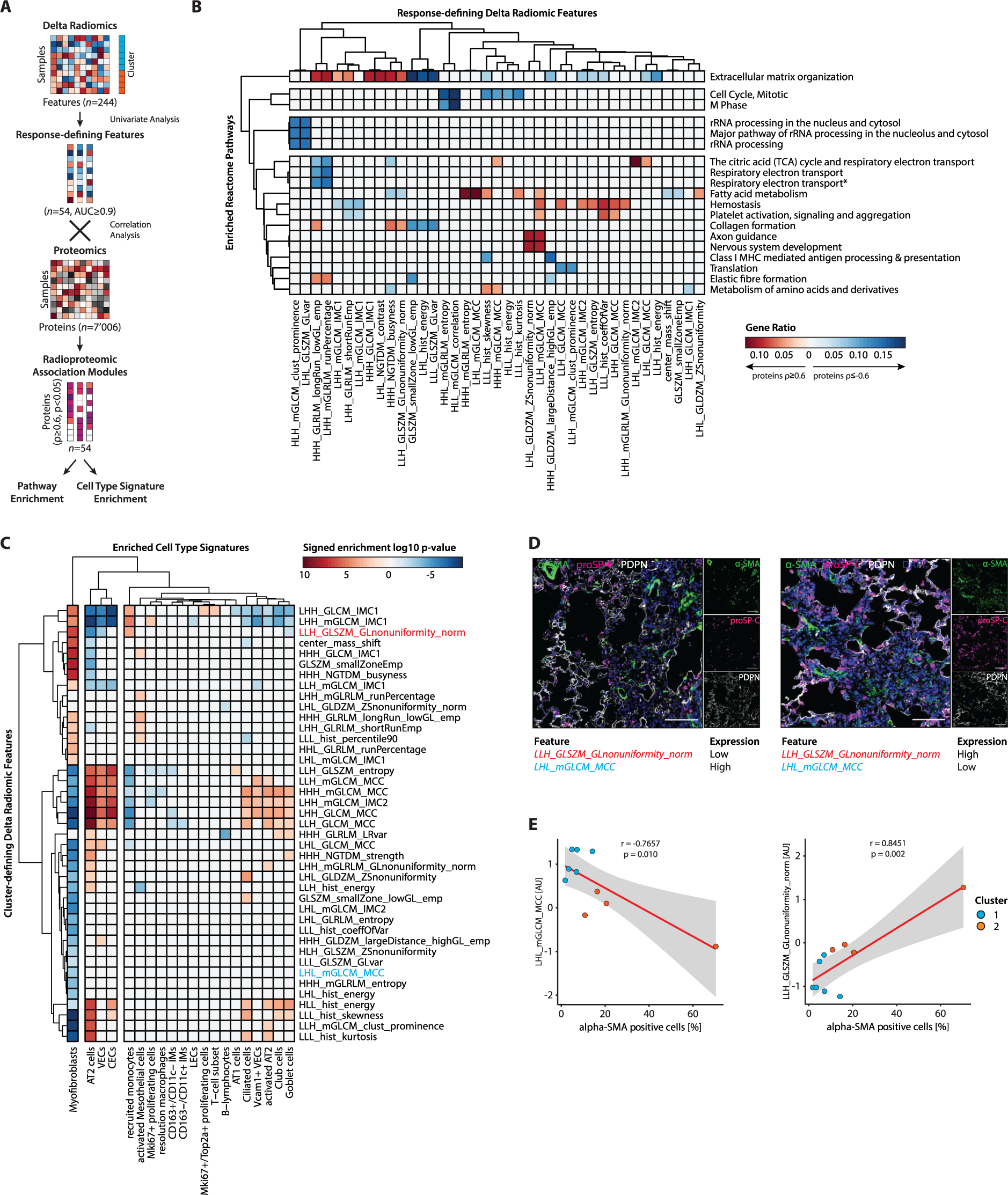
Delta radiomic features reflect changes in disease-relevant molecular pathway activity. (**A**) Schematic of analysis workflow. Variable importance of each delta radiomic feature (*n*=244) for assignment of clusters was assessed by univariate analysis, retaining only “response-defining” features (*n*=54) with classification score≥0.90. Radioproteomic association modules were compiled by assigning the set of highly-correlating proteins (Spearman’s |ρ|≥0.6, p<0.05) to each response-defining feature. These modules were subsequently entered into pathway and cell type signature enrichment analysis. (**B**) Heatmap displaying Reactome pathways enriched (GeneRatio≥0.10, p.adj<0.05) in radioproteomic association modules for positively (Spearman’s ρ≥0.6, p<0.05, red annotation) or negatively (Spearman’s ρ≤-0.6, p<0.05, blue annotation) correlating proteins. Only pathways enrichment in at least two radioproteomic association modules are displayed. Association modules without enriched pathways following filtering are not displayed. (**C**) Heatmap displaying cell type signatures enriched (p<0.01) in radioproteomic association modules for positively (Spearman’s ρ≥0.6, p<0.05, red annotation) or negatively (Spearman’s ρ≤-0.6, p<0.05, blue annotation) correlating proteins. Association modules without enriched cell type signatures following filtering are not displayed. (**D**) Representative IF stainings of fibrotic lung regions exhibiting a low (left) and high (right) fraction of α-SMA+ myofibroblasts. Relative expression of two selected delta radiomic features (*LLH_GLSZM_GLnonuniformity_norm* and *LHL_mGLCM_MCC*) showing positive or negative enrichment for myofibroblast cell type signatures, respectively, is indicated. Images show nuclei (DAPI), AT2 cells (proSP-C), myofibroblasts (α-SMA), and AT1 cells (PDPN). Regions are 500×500 µm in size, scale bar 100 µm. Each data point represents the sample average fraction of α-SMA+ cells of five representative fibrotic regions. (**E**) Scatter plot of the Pearson correlation coefficient between the α-SMA+ cell fraction quantified by IF and the z-scored delta radiomic feature expression of *LHL_mGLCM_MCC* (left) and *LLH_GLSZM_GLnonuniformity_norm* (right). The red line represents the linear model of the best fit, with the gray area representing the 95% confidence intervals. The assigned cluster of each sample is indicated.

To assess if delta radiomic features could provide further insights into changes at the cellular level, we performed cell type deconvolution analysis of the radioproteomic association module-derived protein sets (Spearman’s |ρ|≥0.6, p<0.05) (**Figure 3C, Supplementary Table 13**). Proteins were ranked by log10 p-value and weighted by correlation coefficient prior to entering deconvolution analysis. Overall, we found 41 response-defining delta radiomic features with significant (p<0.01, Kolmogorov-Smirnov test) cell type marker profile enrichment, accounting for 20 different cell types. Noticeably, myofibroblasts, AT2 cells, as well as vascular and capillary endothelial cell gene signatures, showed the most significant correlations. Typically, we observed an inverse correlative relationship between pro-fibrotic and pro-regenerative cell types, as for instance myofibroblasts and AT2 cells. Utilizing immunofluorescence quantification of α-SMA+ myofibroblasts, we validated the top positive and negative correlating features, *LLH_GLSZM_GLnonuniformity_norm* (Pearson’s r=0.85, p=0.002) and *LHL_mGLCM_MCC* (Pearson’s r=-077, p=0.010), which demonstrated significant correlations with myofibroblasts in fibrotic regions (**Figure 3D-E**).

Collectively, our results demonstrated that delta radiomic features capture changes of highly specific molecular and cellular information, thereby highlighting their potential as surrogates for molecular treatment response phenotypes.

### Delta radiomics stratifies nintedanib-treated PF-ILD patients according to lung function decline

We previously demonstrated the high transferability of radiomic signatures from experimental models to human ILD ^37^. To assess whether our preclinical delta radiomic features could stratify nintedanib-treated patients based on their extent of lung function decline, we retrospectively analyzed delta radiomic feature profiles of 19 patients with PF-ILD that received antifibrotic therapy for a median of 12.2±5.7 months (median±IQR) (**Figure 4A**). ILD etiologies included IPF (*n*=11), SSc-ILD (*n*=4), hypersensitivity pneumonitis (HP, *n*=3), and drug-induced ILD (*n*=1) (**Table 1**).

**Figure 4.**
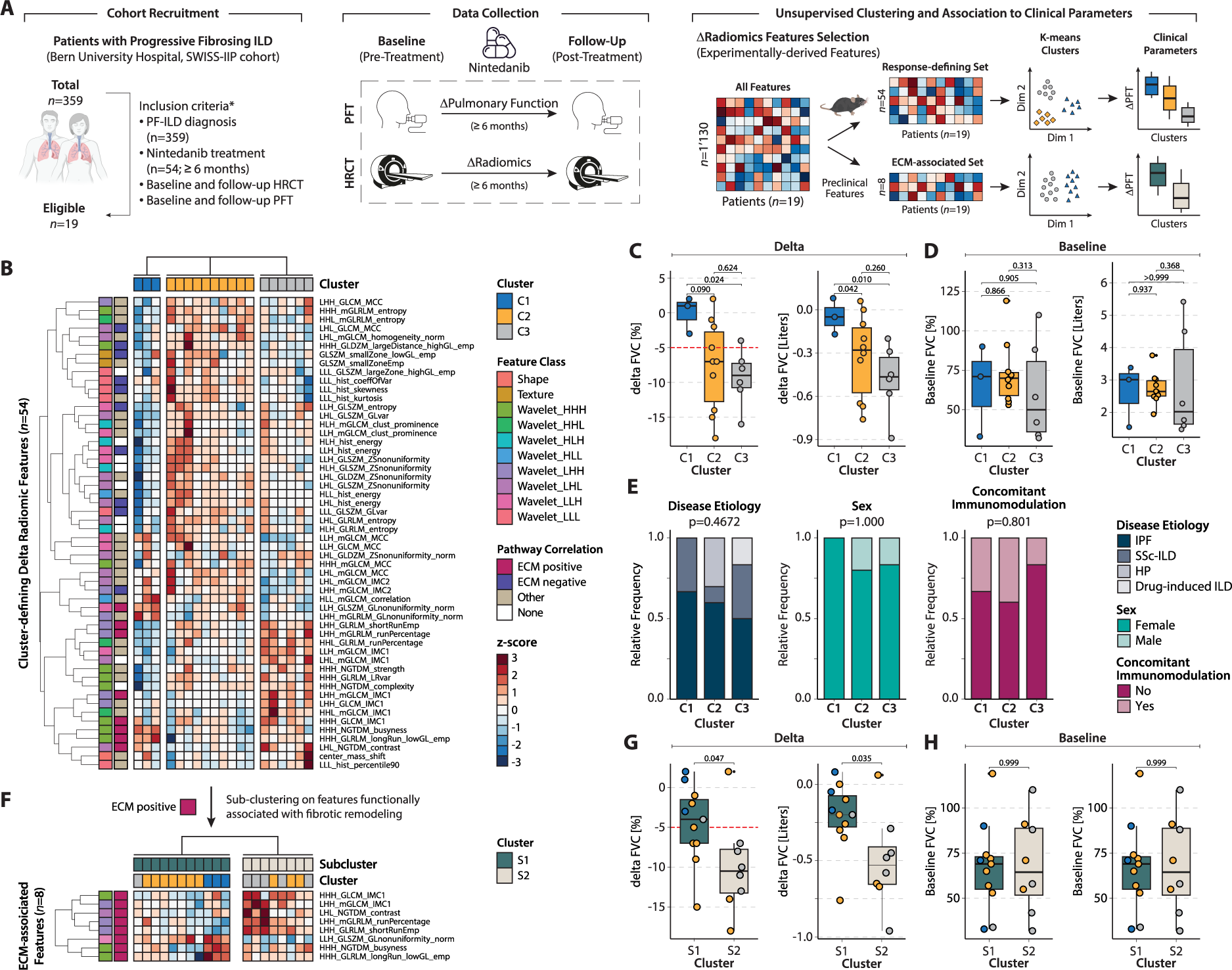
Delta radiomics stratifies nintedanib-treated PF-ILD patients according to lung function decline. (**A**) Workflow schematic. We retrospectively included patients (*n*=19 out of 359 patients) with PF-ILD undergoing treatment with nintedanib at Bern University Hospital and the SWISS-IIP cohort fulfilling the inclusion criteria. For each patient, changes in pulmonary function parameters and radiomic features were calculated between pre- and post-treatment. Unsupervised k-means clustering of patients was performed on subsets of experimentally defined delta radiomic features, including response-defining features (*n*=54) and features positively enriched for ECM remodeling pathway activity (*n*=8). The resulting clusters were investigated for differences in clinical outcome parameters and patient demographics. (**B**) Heatmap displaying the results of unsupervised k-means clustering of the z-scored response-defining delta radiomic feature set (*n*=54) in the PF-ILD cohort. The feature class for each variable and the enrichment of the radioproteomic association module for Reactome pathways is indicated. (**C-D**) Box plots comparing FVC (% pred and liters) delta and baseline level between clusters C1-C3. (**E**) Associations of clusters C1-C3 with clinical and demographic parameters in the PF-ILD cohort. Fisher’s exact test was used to compare the categorical variables. (**F**) Heatmap displaying the results of unsupervised k-means sub-clustering of the z-scored features whose radioproteomic association modules are positively enriched with ECM remodeling Reactome pathway activity (n=8) in the PF-ILD cohort. The feature class for each variable and the enrichment of the radioproteomic association module for Reactome pathways is indicated. (**G-H**) Box plots comparing FVC (% pred and liters) delta and baseline level between clusters S1 and S2.

Unsupervised k-means clustering on the preclinical response-defining delta radiomic feature set (*n*=54) revealed three fairly robust (Jaccard indices>0.60) clusters C1-C3 within the nintedanib-treated PF-ILD cohort (**Figure 4B**, **Supplementary Figure 4A**). These clusters stratified patients according to their annual rate of lung functional decline (**Figure 4C**, **Supplementary Figure 4B, Supplementary Table 14**). Within the observational period, cluster C1 showed a significantly (p<0.05, Mann-Whitney U test) lower FVC decline (1.0±2.5%, -50±125 mL; median±IQR) compared to cluster C3, which showed a substantial decline (-9.0±3.5%, -465±225 mL, median±IQR). Cluster C2 presented with an intermediate phenotype with a considerable FVC decline (-7.0±10.0%, -280±450 mL, median±IQR), although not statistically different from clusters C1 and C3 (p>0.05, Mann-Whitney U-test). Notably, no significant differences (p>0.05, Mann-Whitney U or Fisher’s Exact test) between the clusters were observed for pre-treatment FVC levels (**Figure 4D)**, disease etiology, sex, smoking status, disease duration, presence of concomitant immunomodulatory therapy, or presence of pulmonary (arterial) hypertension (**Figure 4E**, **Supplementary Figure 4C**).

Our preclinical results revealed subsets of radioproteomic association modules that were specifically linked to ECM remodeling activity, the key molecular target of antifibrotic therapy ^35^. To evaluate if the corresponding delta radiomic features (*n*=8, positive enrichment) would lead to improved stratification, we performed k-means clustering of the PF-ILD cohort using this feature subset (**Figure 4F**, **Supplementary Figure 4D**). The resulting two stable clusters S1 and S2 (Jaccard indices>0.75) exhibited significant differences (p<0.05, Mann-Whitney U test) on delta FVC level, with cluster S1 (-4.0±5.5%, -200±205 mL, median±IQR) displaying less functional decline compared to cluster S2 (-10.5±5.5%, -530±245 mL, median±IQR). Notably, this effectively redefined the previous intermediate cluster C2 into either cluster S1 or S2, corresponding to low/intermediate and high FVC decline (**Figure 4G, Supplementary Table 15**). Similar to our previous findings, we found no significant differences between clusters S1 and S2 for pre-treatment FVC levels (p>0.05, Mann-Whitney U-test) (**Figure 4H**) or demographic and clinical variables (p>0.05, Fisher’s exact test) (**Supplementary Figure 4E-F**).

## Discussion

Accurate monitoring of response to antifibrotic therapy is an urgent need for effective management of patients with PF-ILD. However, differentiation between natural disease progression and treatment failure is difficult by means of conventional PFT and HRCT assessment ^6^. Molecular response markers, including peripheral blood biomarkers may improve precision, but these are still in early developmental stages and may not fully reflect lung tissue phenotypes ^7,38,39^. Over the last decade, radiomics has emerged as a powerful tool for drug response monitoring and predicting outcomes in various diseases, such as cancer, neurological disorders and recently also ILDs ^40–42^. The strength of radiomics lies in its ability to provide integrated information on whole lung tissue pathology, conveying both structural and molecular information ^15,17,43^.

In this study, we employed an integrative radioproteomics approach to demonstrate that CT-based delta radiomic profiling can non-invasively stratify the molecular response to nintedanib treatment in a preclinical bleomycin-induced lung fibrosis model, which was not discernible through conventional histogram-based CT measures. We discovered distinct radioproteomic association modules that conveyed disease and drug-specific biological pathway activities and cell type signatures, including ECM remodeling, hemostasis, and fibroblast activation, respectively. Evaluating the preclinical response-defining delta radiomic features, in particular the ECM-associated features in a nintedanib-treated PF-ILD cohort accurately stratified patients according to their extent of lung function decline.

Previous reports have shown the potential of CT-derived imaging characteristics for assessing the response to antifibrotic treatment. Lung attenuation histogram-derived measures, for example, have proven reliable in studying the efficacy of antifibrotic drugs in preclinical lung fibrosis models ^32,44,45^. However, their use as surrogate markers is mostly limited to macroscopic tissue pathologic properties, falling short in resolving the underlying molecular landscape, as also evidenced in the current study. In addition, these variables represent the summary of gray-level intensities not taking the spatial interrelationship of voxels into account. This potentially limits their sensitivity to capture the subtle changes induced by antifibrotic treatment in PF-ILD patients, who often present with morphologically complex and heterogeneous disease patterns ^46,47^. In contrast, higher-order radiomic features, such as texture features quantify the spatial variations in image characteristics, offering added information for treatment monitoring. Utilizing a texture-based quantitative lung fibrosis (QLF) score Kim *et al.* were able to stratify IPF patients undergoing experimental antifibrotic treatment according to the rate of pulmonary function decline ^48^. Furthermore, Devkota *et al.* showed that texture-derived nano-radiomics and not conventional quantitative CT features captured treatment-induced changes of cellular therapy in tumor xenografts ^16^.

The added and complementary value of radiomics arises from the integrated in-depth analysis of tissue heterogeneity across spatial scales conveying pathophysiological information of the whole organ. Imaging omics approaches, including radiogenomics, -transcriptomics, and -proteomics investigate the association between macroscopic radiomic and microscopic molecular features derived from genomic, transcriptomic, or proteomic profiling, respectively to define the underlying biological basis of imaging phenotypes and derive non-invasive imaging surrogates for molecular profiles ^49^. So far, imaging omics have nearly exclusively been studied in the context of cancer. For instance, recent studies utilized radiogenomics to unravel intratumoral heterogeneity phenotypes in multi-center breast cancer cohorts ^18,50^ and identified activated ferroptosis pathways to be associated with high tumor heterogeneity ^18^. Moreover, radiogenomics has been employed to non-invasively characterize the biological activities of specific breast cancer subclones ^50^.

In this study, we provide first evidence that delta radiomic signatures are sensitive towards antifibrotic therapy-induced molecular changes in experimental fibrosing ILD. We add novelty by integrating delta radiomics with proteomics and utilizing the resulting association modules to functionally explain different treatment response phenotypes on a pathophysiologic level. The ability to assess distinct pathway and cellular activities non-invasively from standard-of-care HRCT scans could pave the way towards digital molecular disease fingerprints that could inform precision medicine ^51^.

Our study has some limitations. First, in our preclinical studies, the absence of pre-treatment proteome profiles in mice did not allow us to investigate the molecular landscape at therapy start, which may have confounded the antifibrotic treatment response. However, unlike human disease, inter-individual variance of lung fibrosis development in mice is considered to be low in presence of high bleomycin doses ^52–54^. Future validation of our findings in independent lung fibrosis models will be necessary to ensure the broader applicability of our approach. Secondly, generalizability of our findings to human ILD is limited by the pilot character and retrospective nature of our study. Although we could not find statistically significant differences in potential confounders, we cannot rule out that factors such as concomitant immunomodulatory therapy may have contributed to the effects observed on delta radiomic level given the relatively small sample size. Furthermore, the lack of pre- and post-treatment biosamples precluded molecular validations. Future prospective multi-center studies which include the collection of liquid biopsies for molecular evaluation, together with the inclusion of a placebo group will be necessary to fully elucidate the applicability of delta radiomic signatures as a digital fingerprint for disease or drug-response monitoring. Nonetheless, our ability to detect significant changes in the extent of pulmonary function decline based on preclinical functionally described delta radiomic features in this small but well-defined cohort showcases the method’s inherent potential. Finally, due to the small sample sizes, we could not yet assess the predictive potential of delta radiomic profiles for treatment response, which will be the subject of future studies.

In conclusion, this study highlights delta radiomics as a non-invasive tool to stratify response to antifibrotic treatment in experimental fibrosing ILD through its ability to decode tissue-underlying molecular information. Its potential for transferability to human disease is a first step towards precision medicine, facilitating individual therapy monitoring and risk-benefit assessment in the context of lifelong therapies.

## Author contributions

**DL, JGS** conceived and designed the study, acquired, analyzed and interpreted the data, designed the figures, and wrote the manuscript. **CYM, LK, GMC, AS, MFC, LE** performed the acquisition and analysis of patient data. **HW** performed immunofluorescence stainings. **MB** contributed to the acquisition and analysis of animal and patient data. **MC, HG, STS, KK, OD, SV** provided intellectual input. **AU, MH** contributed to the acquisition and analysis of proteome data. **BM** contributed to the design and conception of the study and wrote the manuscript. All authors read the final manuscript.

## Acknowledgements

We acknowledge Dr. Lutz Wollin (Boehringer Ingelheim Pharma GmbH & Co. KG, Biberach an der Riss, Germany) for supplying us with nintedanib for experimental use and for his advice. We further thank Andrea Laimbacher (Department of Rheumatology, Center of Experimental Rheumatology, University Hospital Zurich, University of Zurich, Zurich, Switzerland) for technical assistance with *in vivo* experimentation and histological analyses. Microscopic imaging was performed with equipment maintained by the Center for Microscopy and Image Analysis, University of Zurich, Zurich, Switzerland. Proteome analyses were performed on mass spectrometry equipment financed by the Swiss National Science Foundation (R’Equip grant No. 316030-189737) and the University of Bern at the Proteomics & Mass Spectrometry Core Facility, University of Bern, Bern, Switzerland.

## Data availability

All data and code for reproduction of the main findings of this study will be made publicly available after publication.

## Supplementary Figures

**Supplementary Figure 1.**
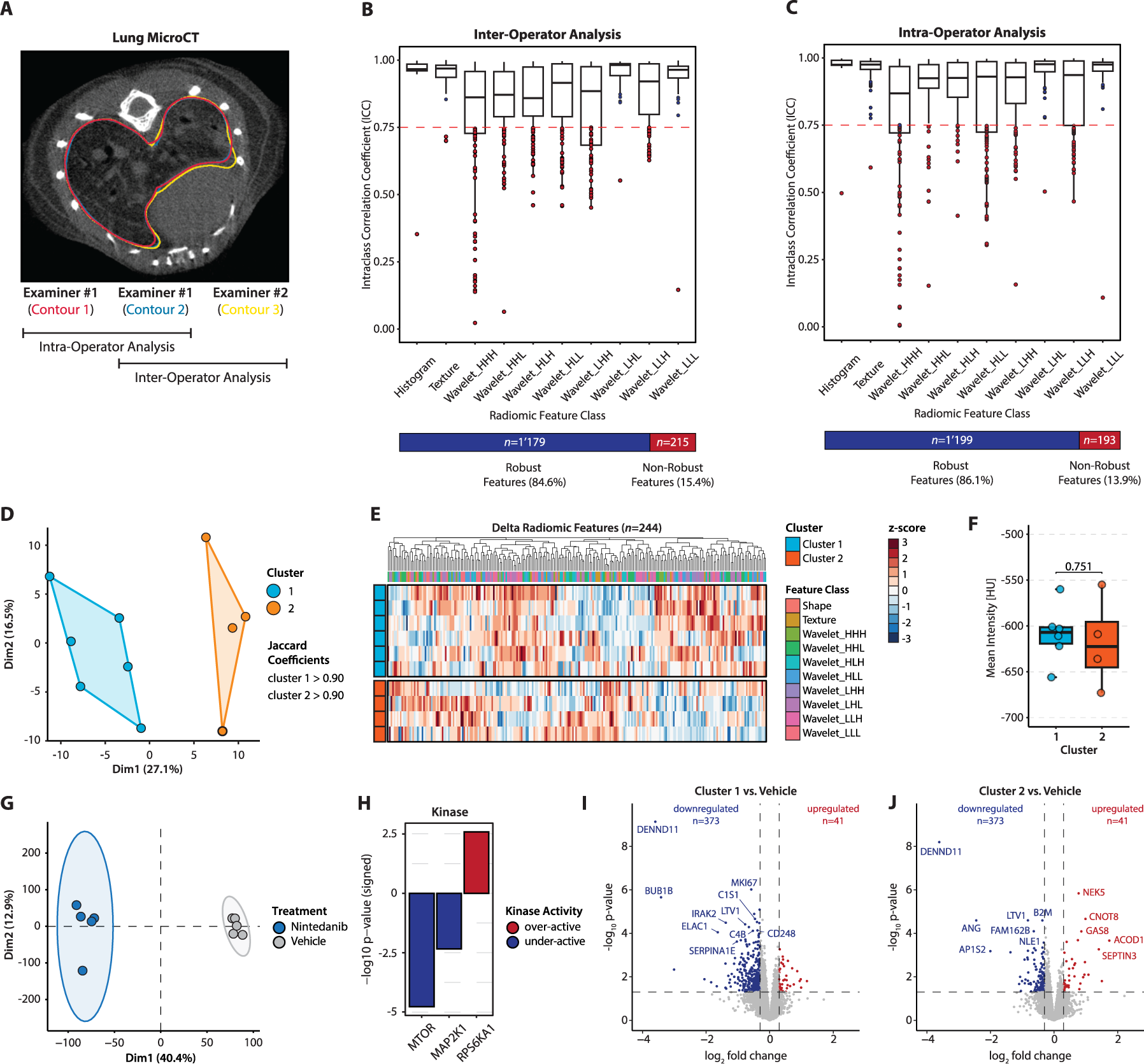
(**A**) Evaluation of the radiomic feature stability against inter- and intra-reader variation in the semi-automated lung segmentation workflow. A representative transversal microCT image of a bleomycin-induced mouse lung is displayed. Outlined are three semi-automatically delineated lung contours of two different examiners (examiner 1: red and blue; examiner 2: yellow). For intra- and inter-operator ICC analysis, a total of *n*=16 randomly selected lung scans covering different time points were segmented in this manner. (**B**) Boxplots displaying the distribution of the ICC coefficient per radiomic feature class for inter-operator ICC analysis and (**C**) intra-operator ICC analysis. The red dashed line indicates the set intraclass correlation coefficient (ICC) threshold at 0.75. The stacked bar charts summarize the relative frequency and total number of robust and non-robust radiomic features. (**D**) K-means sub-clustering of z-scored delta radiomic features (*n*=244) of nintedanib-treated mice (*n*=10) indicates two stable clusters (Jaccard coefficients>0.90, where 1 describes perfect stability). (**E**) Heatmap summary of the k-means sub-clustering results (nintedanib-treated mice, *n*=10). Clusters and the feature class of each variable are indicated. (**F**) Lung tissue density in cluster 1 and 2 expressed as mean Hounsfield unit (HU) intensity post-treatment. Mann-Whitney U-test was used to compare the numerical variables. (**G**) Principal component analysis of the phosphosite expression (*n*=20’043) profiles in subsets of randomly selected nintedanib- (*n*=5) and vehicle-treated (*n*=5) mice. (**H**) Kinase activity enrichment analysis (KAEA) of differentially expressed phosphosites in nintedanib against vehicle-treated mice. Under-active kinases colored in red, over-active kinases colored in blue. (**I**) Volcano plots of protein expression in cluster 1 and (**J**) cluster 2 compared to vehicle-treated mice. Proteins with log2FC>0.30 and p<0.05 were considered to be differentially expressed. Down- and upregulated proteins are highlighted in blue and red, respectively.

**Supplementary Figure 2.**
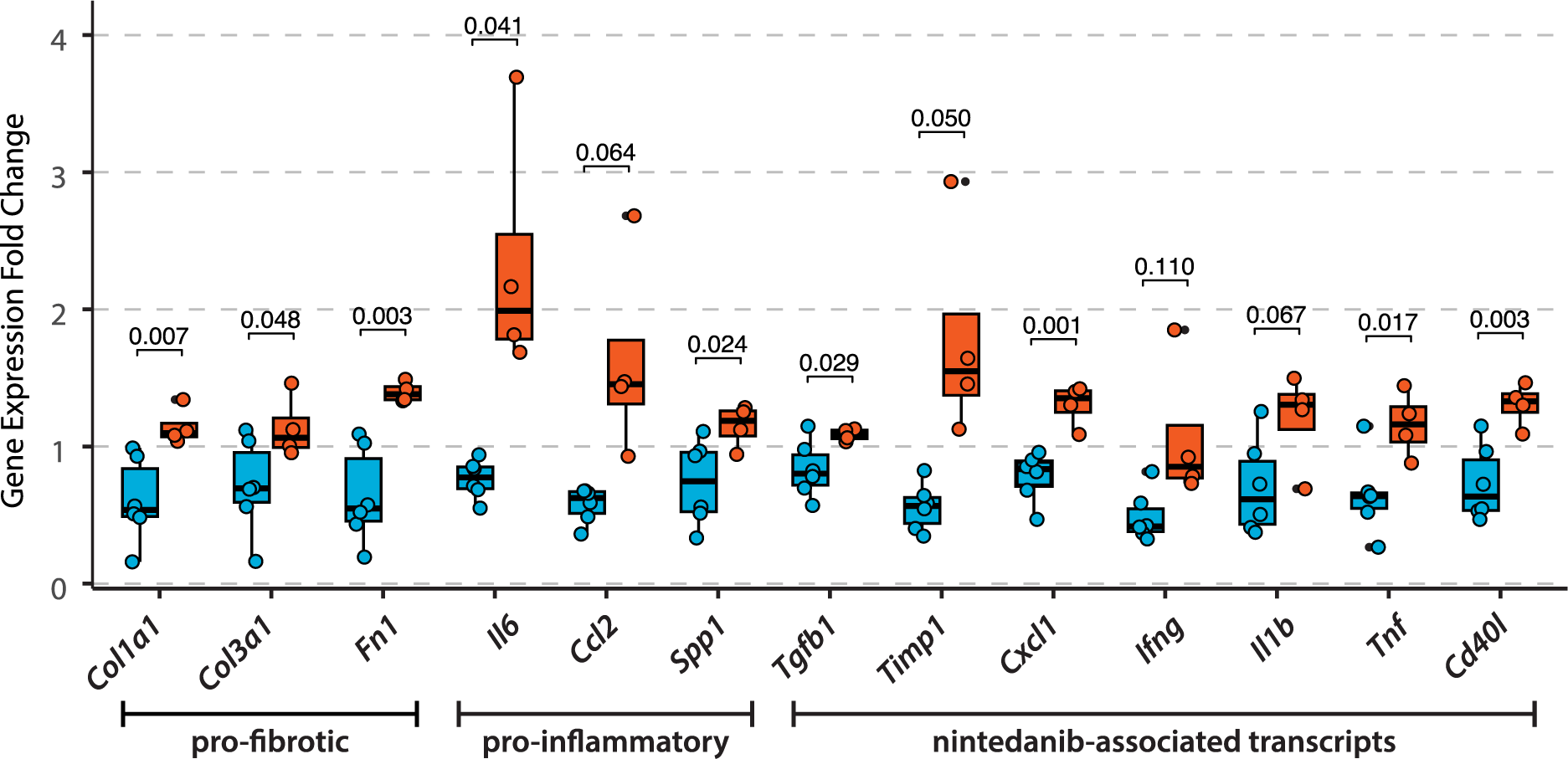
Quantitative PCR of pro-fibrotic (*Col1a1*, *Col3a1*, *Fn1*), pro-inflammatory (*Il6*, *Ccl2*, *Spp1*), and nintedanib-targeted (*Tgfb1*, *Timp1*, *Cxcl1*, *Ifng*, *Il1b*, *Tnf*, *Cd40l*) genes. Displayed is the mRNA fold change expression using the 2^-ΔΔCt^ method in cluster 1 (blue) and cluster 2 (red) over vehicle-treated samples. Each data point represents the mean of two technical replicates. Unpaired Student’s t-test was used to compare the continuous variables.

**Supplementary Figure 3.**
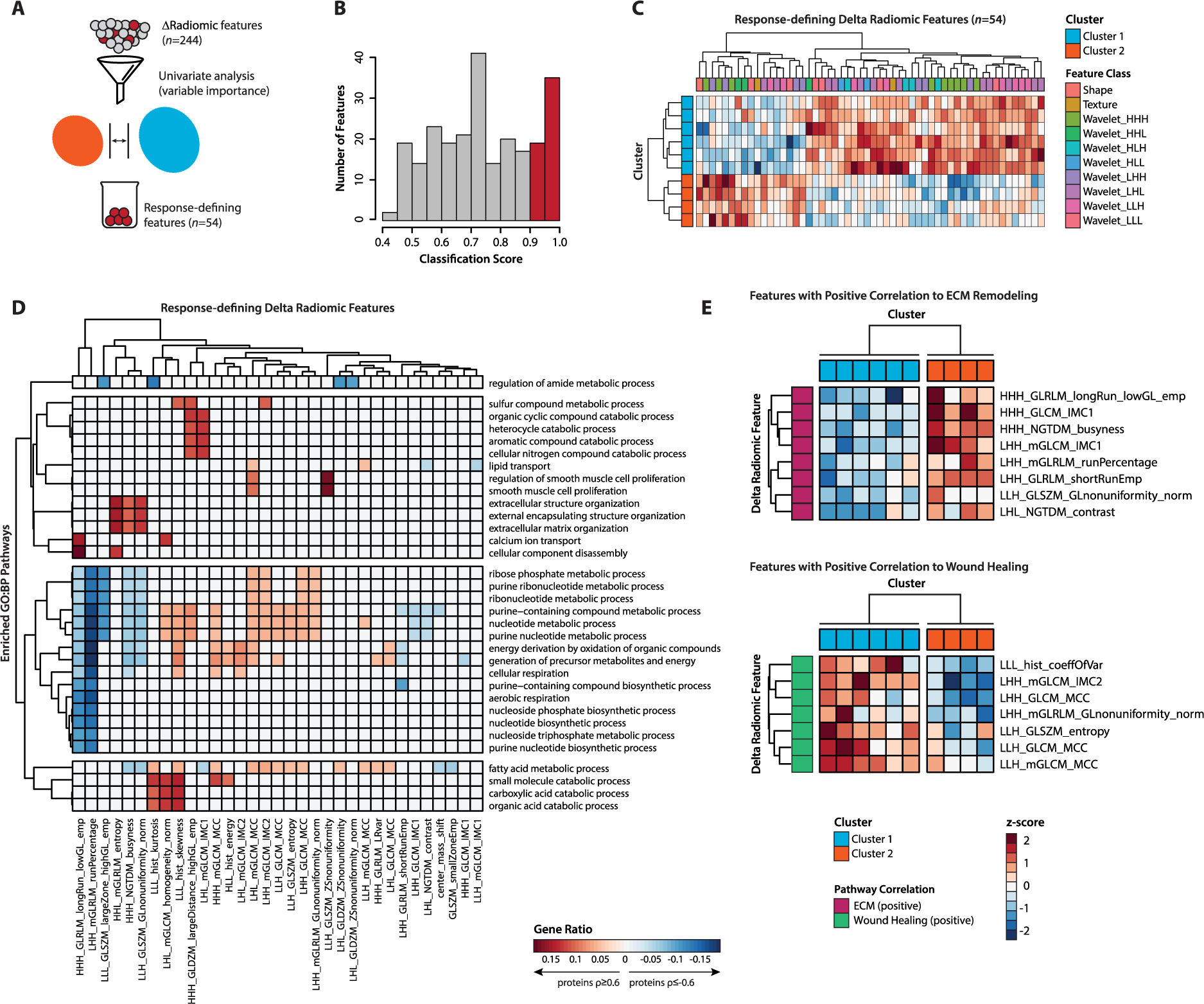
Identification of response-defining delta radiomic features and their correlation with disease-relevant pathways. (**A**) Variable importance of each delta radiomic feature (*n*=244) for assignment of clusters was assessed by univariate analysis, retaining only “response-defining” features (*n*=54) with classification score≥0.90. (**B**) Histogram of the univariate analysis results of the delta radiomic features (*n*=244) for classification of cluster assignment. Variables with classification score≥0.90 (red) were considered to have response-defining properties. (**C**) Heatmap displaying the results of unsupervised hierarchical clustering of z-scored subset of response-defining delta radiomic features (*n*=54) in nintedanib-treated mice (*n*=10). Cluster assignment and the feature class of each variable are indicated. (**D**) Heatmap displaying GO:BP pathways enriched (GeneRatio≥0.10, p-adjusted<0.05) in radioproteomic association modules for positively (Spearman’s ρ≥0.6, p<0.05, red annotation) or negatively (Spearman’s ρ≤-0.6, p<0.05, blue annotation) correlating proteins. Only pathways enriched in at least two radioproteomic association modules are displayed. Association modules without enriched pathways following filtering are not displayed. (**E**) Heatmaps displaying the results of unsupervised k-means clustering of z-scored subsets of delta radiomic features positively enriched in ECM remodeling (*n*=8) and wound healing (*n*=7) in nintedanib-treated mice (*n*=10), respectively. Cluster assignment of samples and Reactome pathway enrichment of variables are indicated.

**Supplementary Figure 4.**
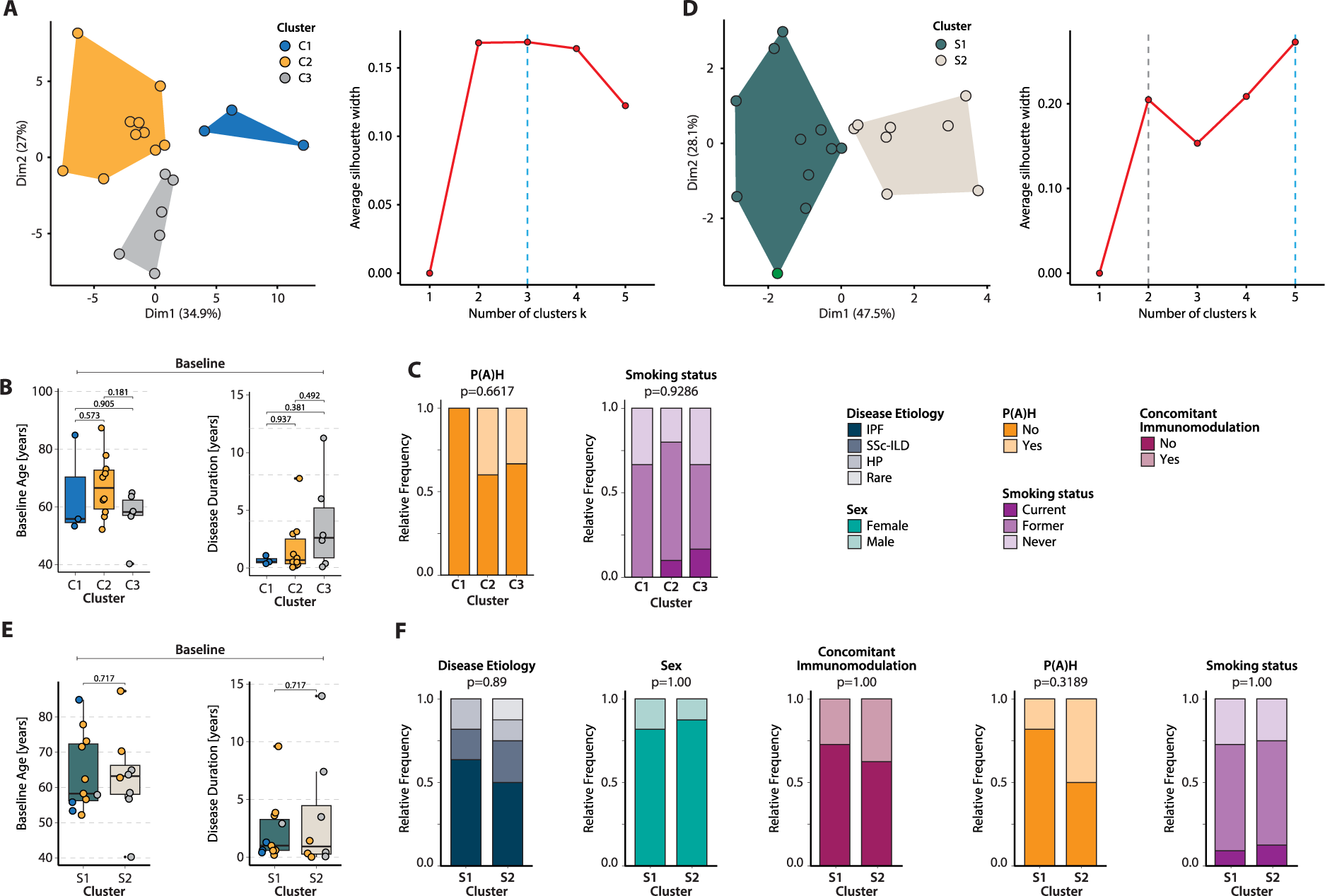
Delta radiomics stratifies the degree of pulmonary function decline in nintedanib-treated PF-ILD patients. (**A**) Left: K-means cluster plot for z-scored preclinical response-defining delta radiomic features (*n*=54) of the PF-ILD cohort (*n*=19) indicates three fairly robust clusters (Jaccard coefficients>0.60, where 1 describes perfect stability). Right: Scatter plot showing the average silhouette coefficient versus the number of clusters for the k-means clustering input data. The blue dashed line indicates the global optimum. (**B**) Box plots comparing age (years)^#^ and disease duration (years)^¶^ at baseline between clusters C1-C3. Mann-Whitney U test was used to compare the continuous outcomes. (**C**) Associations of clusters C1-C3 with clinical and demographic parameters in the PF-ILD cohort. Fisher’s exact test was used to compare the categorical variables. (**D**) Left: K-means cluster plot for z-scored delta radiomic features positively correlating with ECM-remodeling (*n*=8) of the PF-ILD cohort (n=19) indicates two stable clusters (Jaccard coefficients>0.75, where 1 describes perfect stability). Right: Scatter plot showing the average silhouette coefficient versus the number of clusters for the k-means clustering input data. The blue and red dashed line indicate the global and local optimum, respectively. (**E**) Box plots comparing age (years)^#^ and disease duration (years)^¶^ at baseline between clusters S1 and S2. Mann-Whitney U test was used to compare the continuous outcomes. (**F**) Associations of clusters S1 and S2 with clinical and demographic parameters in the PF-ILD cohort. Fisher’s exact test was used to compare the categorical variables. ^#^Age is defined as the period between birth date and baseline HRCT scan. ^¶^Disease duration is defined as the period between the first reported diagnosis of PF-ILD and baseline HRCT scan.

## Supplementary Methods

### Animal experimentation and ethics statement

The effects of antifibrotic treatment on delta radiomics were studied in mice with bleomycin-induced lung fibrosis ^22,23,32,55^. To induce lung fibrosis, C57BL/6J-Rj mice (*n*=30, female, 8-weeks old, Janvier Labs, Le Genest-Saint-Isle, France) were intratracheally instilled with 2 U/kg bleomycin sulfate (Bleomycin Baxter 15’000 I.U., pharmacy of the canton Zurich, Switzerland) dissolved in saline on day 0. Treatment with 60 mg/kg nintedanib (*n*=15) or vehicle-only (deionized water, *n*=15) was administered once daily by oral gavage (at a volume of 10 µL/g body weight) from day 7 to day 20 for a total of 14 applications. Lung MicroCT scans were acquired of each animal pre- (day 7) and post-treatment (day 21) for generation of radiomic feature sets. All mice were sacrificed on day 21 by CO_2_ inhalation (24 hours after the final treatment), followed by exsanguination of the vena cava and transcardial perfusion of the lungs with 10 mL ice-cold Dulbecco’s phosphate-buffered saline (DPBS) at a pressure of 100-120 cm H_2_O to remove residual blood from the lung. The lung was excised, rinsed with DPBS, dissected into the individual lobes, and processed according to the different assay requirements. Mice were allocated to different study groups by complete randomization and treatment was provided in a double-blinded manner. All mice were housed in groups of five with access to food and water *ad libitum* in standard housing conditions with 12 hour light-dark cycles. Animals were acclimatized for seven days prior to experimentation start. HydroGel® (ClearH2O Inc.) and water-soaked standard rodent diet were provided to alleviate body weight loss. Paracetamol analgesia was provided if mice showed signs of pain. Ethical approval for experimentation was granted by the cantonal veterinary office (license no. ZH082/2021) and experimentation was performed in strict compliance with Swiss animal protection laws and guidelines. Mice were excluded from analyses if humane endpoints were reached (*n*=3) or if microCT scans exhibited presence of severe lung abnormalities (*n*=3), including atelectasis or unilateral fibrosis development.

### Patient cohort, clinical data, and ethics statement

In this study, 19 PF-ILD patients undergoing treatment with nintedanib at Bern University Hospital were retrospectively selected from the Bern University Hospital registry and the SWISS-IIP cohort. Approval for the study was granted by the local ethics committee (BASEC-ID: 2023-01920 [ILDALMO]; PB_2016_01524 [SWISS-IIP cohort]). Selection was performed based on the inclusion and exclusion criteria stated below. A total of 359 patients diagnosed with (progressive) fibrosing ILD were screened for fulfillment of the below criteria, of which 54 patients received nintedanib treatment for ≥6 months. Of these 54 patients, 19 fulfilled also the remaining inclusion criteria.

### Inclusion criteria

1. Diagnosis of PF-ILD determined by a senior attending physician according to established guidelines ^2,24^
2. Treatment with nintedanib (≥100 mg twice daily; min. 6 months at follow-up HRCT)
3. Availability of HRCT chest scans fulfilling the following criteria:

a. Pre-treatment HRCT (max. 1 month after treatment initiation)
b. Post-treatment HRCT (min. 6 months interval to pre-treatment HRCT)
c. Slice thickness in range 0.5 - 1.5 mm
d. Acquisition at tube voltage in range 80-130 kVp
e. One of the following reconstruction kernels: I70f, I80s, Br56f, Br56u, Br59f, LUNG, FC55 (sharp), YB
f. Filtered-back projection as reconstruction algorithm
g. Scans acquired in full inspiration mode
4. Availability of PFT recordings fulfilling the following criteria:

a. Pre-treatment PFT (max. 1 month after treatment initiation)
b. Post-treatment PFT (min. 6 months interval to pre-treatment PFT)

### Exclusion Criteria

5. Presence of secondary lung disease at times any HRCT scans and PFT recording (e.g., cancer, COVID-19, pneumonia, bronchitis)
6. Concomitant treatment with other antifibrotic drugs during the observation period (e.g. pirfenidone)

Demographic and clinical parameters were derived from electronic patient records, including age (birth date), sex, disease etiology, date of diagnosis, date of nintedanib treatment start, presence of pulmonary (arterial) hypertension, smoking status, concomitant medications, and dates of PFT and HRCT scan recordings. The recorded PFT parameters included forced vital capacity (FVC) in % pred. and liters. Changes in PFT recordings between pre- and post-treatment were expressed as delta values. A Summary of patient demographics and clinical characteristics is provided in **Table 1**.

### MicroCT image acquisition

Lung microCT images of each mouse were acquired at days 7 and 21 on a SkyScan 1176 (Bruker, Kontich, Belgium) in free-breathing conditions under isoflurane anesthesia using respiratory gated image acquisition. Anesthesia was induced by 5.0% and maintained by 1.5-2.5% isoflurane in air at 0.8-1.0 L/min flow rate to achieve a breathing rate of 0.7-0.9 breaths/s for an average scan time of 15 min. Animals were placed in supine position on the scanner bed with a styrofoam block mounted on the diaphragm to allow monitoring of respiratory gating. Image acquisition was performed with the following acquisition settings: tube voltage = 50 kV, tube current = 500 µA, filter = Al 0.5 mm, frame averaging = on (3), rotation step = 0.7 degrees, sync with events = 50 ms, X-ray tube rotation = 360 degrees, exposure time = 77 ms, resolution = 35 µm, slice thickness = 35 µm. Images were reconstructed with NRecon software (v.1.6.8.0; Bruker) using Feldkamp filtered back-projection algorithm with the following parameters: misalignment compensation (scan-dependent manual adjustment), smoothing = 1 with Gaussian kernel, ring artifact compensation = 4, and beam hardening correction = 10%. Reconstructed images were converted to DICOM format.

### CT segmentation of mouse lungs

Left and right lung lobes of mice were semi-automatically segmented by two readers (D.L., M.B.) using MIM software (v.7.1.6, MIM Software Inc., Cleveland, Ohio, USA). Briefly, a seed was set within the right and left lung using the “region grow” tool (upper limit = -600 HU, lower limit = -800 HU, tendril diameter = 0.2 mm, fill holes = strong), which then automatically defined the vast majority of the lung volumes in the 3D space. Manual contour alignment with the 2D/3D brush was used to correct potentially misaligned areas. Finally, the smoothing function was used to remove sharp edges. For medical diagnostics, the Hounsfield scale is usually normalized to 120 keV tube voltage, which could technically not be achieved by our microCT instrument. To enable direct comparison between CT-derived radiomic datasets from patients and mice, the reconstructed microCT images of mouse lungs were pixel value corrected to match clinical specifications as previously described ^17^.

### HRCT image acquisition

HRCT acquisition of human lungs was performed at Bern University Hospital or outpatient clinics. Instrument and scan settings used are summarized in **Supplementary Table 1**. All HRCT scans were evaluated by a senior radiologist (L.E.) at the Department of Diagnostic, Interventional, and Pediatric Radiology of the Bern University Hospital for the presence of PF-ILD on a standard picture archiving and communication system workstation and a radiology-grade display monitor.

### CT segmentation of human lungs

Left and right lung lobes were semi-automatically segmented by two readers (C.M., L.K.) with the open source software 3D Slicer (v.5.2.1). Pulmonary hilar vessels and atelectatic areas were manually excluded from the regions of interest. Manual contour corrections were only applied when spatially limited areas did not coincide with the actual borders of the lungs.

### Pulmonary function tests

Pulmonary function tests were performed by trained personnel at the Department of Pulmonary Medicine of the Bern University Hospital or in outpatient clinics. All tests were performed following established protocols ^56–59^.

### Radiomic feature calculation

Calculation of radiomic features was performed on merged structures of left and right lung lobes using Z-Rad software (v.7.3.1, https://medical-physics-usz.github.io/, Department of Medical Physics, University Hospital Zurich, Zurich, Switzerland), an image biomarker standardization initiative (IBSI)-compliant Python-based software ^60^, as described in ^17^. Mouse lungs were resized to isotropic voxels of 0.15 mm. To achieve comparable voxel size in patients, human lungs were resized to isotropic voxels of 2.75 mm, corresponding to an estimated 6000-fold volumetric difference ^26^. Both mouse and human lung volumes were discretized to a fixed bin size of 50 HU in a range of -1000 HU to 200 HU. From the resized volumes, 1’388 radiomic features were calculated per lung scan and time point (HU limits: -1000 to 200 HU), corresponding to the following feature classes:

1. Histogram features (*n*=17)
2. Texture features (*n*=137): Gray Level Co-occurrence Matrix (*n*=52, GLCM), Neighborhood Gray Tone Difference Matrix (*n*=5, NGTDM), Gray Level Run Length Matrix (*n*=32, GLRLM), Gray Level Size Zone Matrix (*n*=16, GLSZM), Gray Level Distance Matrix (*n*=16, GLDZM), and Neighboring Gray Level Dependence Matrix (*n*=16, NGLDM)
3. Wavelet features (*n*=1’232): Transformation of histogram and texture features following coiflet filter decomposition
4. Shape features (*n*=2)

Histogram features carry information about distribution of voxel intensities using first-order statistics (e.g. mean, standard deviation, skewness, kurtosis), describing tissue intensity characteristics. Texture features define intra-tissue heterogeneity by calculating the spatial relationship between neighboring voxel intensities ^61^. Wavelet features compute histogram and texture features after wavelet decompositions of the original image using eight different coiflet filters (high- to low-pass filters), thereby concentrating the features on different frequency ranges ^62^. Shape features describe tissue volume and size independent of intensity distribution.

Delta radiomic features describing the change of each feature between pre-and post-treatment, were expressed as delta values: ΔFeature = Feature (t_2_) – Feature (t_1_) ^27^. Lung densitometric information was directly inferred from the radiomic histogram feature *hist_mean*, which describes the lung attenuation-based average HU intensity of the segmented lung volume.

### Radiomic feature stability evaluation

Intraclass correlation coefficients (ICC) were calculated for each radiomic feature to evaluate stability against inter- and intra-reader bias in the lung segmentation process. For inter- and intra-reader ICC, two (D.L., M.B.) and one examiner(s) (D.L.), respectively, independently segmented 16 randomly selected mouse lung scans, followed by radiomic feature calculation of the delineation structures. ICCs were calculated using two-way mixed effect models with the *consistency* method in the “irr” R package according to published reports ^63^. Only stable/reproducible features (*n*=1’130) with ICC≥0.75 were considered for further analyses for both mouse and human datasets ^64^. Feature stability assessment was performed on the mouse dataset due to the lesser degree of automation in lung segmentation.

### Proteomics

For comparative proteomics, the middle lobe of the right mouse lung was snap frozen in liquid nitrogen and stored at -80°C until processing. Sample workup and data collection was performed by trained personnel at the Proteomics and Mass Spectrometry Core Facility (PMSCF) at the University of Bern using standard established protocols. All vehicle- (*n*=14) and nintedanib-treated (*n*=10) samples were analyzed. One vehicle sample was excluded from analysis due sample workup issues. Tissue homogenization was performed in 8M urea / 100 mM Tris (pH 8.0) buffer supplemented with *cOmplete* protease inhibitor cocktail (Roche Diagnostics, Mannheim, Germany) using the FastPrep system (MP Biomedicals). Following reduction, alkylation, and overnight protein precipitation with ice-cold acetone, 10 µg of the cleaned protein mixture was digested into peptides using a two-step digestion protocol (LysC for 2 h at 37 °C followed by Trypsin at room temperature overnight). Digests were analyzed by nano-liquid chromatography on a Dionex Ultimate 3000 (ThermoFisher Scientific, Reinach, Switzerland) through a CaptiveSpray source (Bruker, Bremen, Germany) with an end-plate offset of 500 V, a drying temperature of 200 °C, and with the capillary voltage fixed at 1.6 kV. A volume of 2 µL (200 ng) protein digest was loaded onto a pre-column (PepMap 100 C18, 5 µm, 100 A, 300 µm diameter x 5 mm length, ThermoFisher) at a flow rate of 10 µL/min with 0.05% trifluoroacetic acid in water / acetonitrile 98:2. After loading, peptides were eluted in back flush mode onto a in-house made C18 CSH Waters column (1.7 µm, 130 Å, 75 µm x 20 cm) by applying a 90-minute gradient of 5% acetonitrile to 40% in water / 0.1% formic acid, at a flow rate of 250 nL/min. The timsTOF Pro instrument (Bruker, Bremen, Germany) was operated either in data-dependent acquisition (DDA) or data-independent (DIA) mode using the Parallel Acquisition Serial Fragmentation (PASEF) option. The mass range was set between 100 and 1700 m/z, with 10 PASEF scans between 0.7 and 1.4 V s/cm^2^. The accumulation time was set to 2 ms, and the ramp time was set to 100 ms, respectively. Fragmentation was triggered at 20’000 arbitrary units, and peptides (up to charge of 5) were fragmented using collision induced dissociation with a spread between 20 and 59 eV. DDA data was processed further with FragPipe software (v.17.0) using the IonQuant algorithm and filtering protein identifications to a 1% false discovery rate (FDR) on the peptide level using the Percolator algorithm. Furthermore, protein groups were filtered by the criterion that at least two different razor peptide sequences were identified as evidence for the existence of the protein group. From the DDA data, a spectral library was built with the FragPipe software. This library was used to identify and quantify proteins with the DIA data using standard parameters in Spectronaut 16 software (Biognosys, Schlieren, Switzerland). Protein names (Uniprot IDs) were converted to Entrez IDs and Gene Symbols using “uniprot.ws” and “annotationDbi” R packages. Protein names without matching Entrez Gene ID were dropped, resulting in a final set of 7’006 proteins.

### Phosphoproteomics

For phosphoproteomics, the middle lobe of the right mouse lung was snap frozen in liquid nitrogen after collection and stored at -80°C until processing. Sample workup and data pre-processing was performed by trained personnel at the Proteomics and Mass Spectrometry Core Facility (PMSCF) at the University of Bern using standard established protocols. Randomly selected subsets of vehicle- (*n*=5) and nintedanib-treated (*n*=5) were analyzed. A titanium dioxide phosphopeptide enrichment workflow ^28^ with subsequent DDA liquid chromatography tandem mass spectrometry (LC-MS) analysis on the same instrument and parameter settings as described above was applied. Samples were searched and quantified with FragPipe ^65^ (v.18.0, MSFragger version 3.5, Philosopher version 4.4.0, IonQuant version 1.8.0) using the following parameters: swissprot ^66^ Mus musculus database (release 2022_01) with isoforms and common contaminants; 20 ppm and 0.05 Da mass tolerance for precursors and fragment, respectively; search enzyme trypsin with max 3 allowed missed cleavages; fix modification: carbamidomethylation of cysteine; variable modifications (altogether max 4/peptide): methionine oxidation of methionine (max 3/peptide), phosphorylation of serine, threonine and tyrosine (max 3/peptide) and protein N-terminal acetylation. Peptide forms normalized with the variance stabilization ^67^ normalization method are reported as NormI, along FragPipe’s MaxLFQ and IonQuant’s FragI abundance measures. The intensities of peptide forms were combined as protein phosphosite locations by summing the corresponding contributions.

### Kinase activity enrichment analysis

Differential expression of phosphosites and subsequent kinase activity enrichment analysis was performed as described in ^28^. First, the phosphosites’ missing values were imputed using a left-censored Gaussian replacement method if there was more than 1 missing value in a group of replicate, and a maximum likelihood estimation otherwise ^68^. A moderated t-statistic ^68^ was then calculated for each phosphosite, and used as the ranking metric for the Kinase Activity Enrichment Analysis (KAEA) tool ^28^. KAEA was then applied on the ranked phosphosite list and reversed ranked list using the included mouse kinase substrate database. SetRank set p-value and FDR cutoff were set to 0.01 and 0.05, respectively.

### Differential protein expression analysis

Differential expression of proteins between groups of interest was calculated in R using the “limma” package according to standard guidelines ^69^. At first, DIA-based Spectronaut protein expression intensities were log_2_-transformed. Then, log_2_ fold changes were calculated as contrasts by application of a linear model using robust regression for each protein. Finally, estimated coefficients and standard errors for the given set of contrasts were calculated for each protein, followed by Empirical Bayes smoothing of standard errors. Proteins with log_2_FC>0.3 (p<0.05), corresponding to 23% mean expression change, were considered as statistically significant.

### Gene expression analysis

RNA was isolated from blood-free cranial lobes of the right mouse lung stored in RNAlater (ThermoFisher Scientific). Tissues were mechanically homogenized with the TissueLyser II instrument (Qiagen, Hombrechtikon, Switzerland), followed by total RNA isolation with the RNeasy Tissue Mini Kit (Qiagen, Hombrechtikon, Switzerland). Isolated RNA was reverse-transcribed into cDNA using the Transcriptor First Strand cDNA Synthesis Kit (Roche Diagnostics, Switzerland). Expression of fibrotic (*Col1a1*, *Col3a1*, *Fn1*), inflammatory (*Ccl2*, *Il6*, *Spp1*), and nintedanib-related (*Tgfb1*, *Timp1*, *Tnf*, *Cxcl1*, *Cd40l*, *Il1b*) genes was analyzed by SYBR Green quantitative PCR using GoTaq Green Master Mix kit (Promega) as described in ^29^. Expression of mRNA was expressed to delta Ct values (Ct [gene of interest] – Ct [reference gene]) with *Rplp0* as reference gene. Lower delta Ct values indicate higher target gene expression. Fold changes relative to vehicle-treated samples were calculated using the delta-delta Ct method. The list of the primer pairs used in this study is provided in **Supplementary Table 2**.

### Immunofluorescence and microscopy

Formalin-fixed paraffin-embedded lung sections (3 µm thickness) were cut on a HistoCore Multicut microtome (Biosystems Switzerland AG, Muttenz, Switzerland). Following deparaffinization, heat-mediated antigen retrieval with R-Universal Buffer (Cat. AP0530-500, Aptum Biologics) was performed for 15 min at 95°C. After incubation for 25 min at RT for cooling, blocking of unspecific antibody staining was performed with 5% BSA in antibody diluent (Cat. S3022, Dako) for 1 h at RT. Primary antibodies dissolved in antibody diluent (Cat. S3022, Dako) were then applied and incubated overnight at 4°C. Next, samples were incubated with secondary antibodies dissolved in PBS supplemented with 1% BSA for 2 h at RT. All antibodies and the dilutions used are listed in **Supplementary Table 3**. Finally, cell nuclei were counterstained with 4’,6-diamidino-2-phenylindole (DAPI) for 10 min at RT. The sections were then scanned in immunofluorescence mode on a AxioScan.Z1 slide scanner (Zeiss, Feldbach, Switzerland) using a Plan-Apochromat 20×/0.8 M27 objective. Cells positively stained for α-SMA were quantified using the “Positive cell detection” tool of the open source software QuPath (v.0.4.0) at default parameter settings and detection thresholds 800, respectively ^70^. From each sample, five representative areas at 500×500 µm were quantified and the average was used for statistical analyses.

### Unsupervised clustering

All variables were z-scored ([x-mean]/standard deviation) prior to analysis (“clusterSim” R package). Unsupervised agglomerative hierarchical (using Euclidean distance with complete linkage method) or k-means clustering was performed to identify subgroups of mice or patients with similar delta radiomic feature patterns or proteomic profiles using base R functions. Clusterability was evaluated by Hopkin’s statistic H, with H>0.5 indicating clusterability (“hopkins” R package) ^71^. The optimal number of clusters was determined by average silhouette statistics inspecting k clusters between 2 and 5, selecting the optimal k based on global or local optimum for separation (“factoextra” R package). Stability of clusters was assessed by Jaccard bootstrapping with n=1’000 iterations (“fpc” R package) ^72^.

### Variable importance evaluation

To estimate the importance of each delta radiomic feature for the classification produced by unsupervised clustering, we calculated filter-based variable importance using the “caret” package ^73^ and retained features with classification score≥0.9. Features (*n*=54) most important for differentiating clusters 1 and 2 in nintedanib-treated mice are listed in **Supplementary Table 4**.

### Gene Ontology and Reactome Pathway enrichment

Curated lists of DE or highly-correlated proteins were used to perform Gene Ontology (GO) or Reactome pathway enrichment analysis using the R packages “clusterProfiler” and “ReactomePA”, respectively ^74–76^, retaining results after false discovery rate adjustment (p<0.05). In case of enrichment of proteins highly-correlated with delta radiomic features, proteins with positive and negative correlation coefficients were entered separately into GO or Reactome pathway enrichment analysis. To visualize and interpret results, the GeneRatios of positive and negative enriched pathways were transformed into matrices with delta radiomic features as columns and pathways terms as rows. Then, results with GeneRatios<0.10 were dropped (set to zero), and only pathway terms enriched (GeneRatio≥0.10) in at least two delta radiomic features were retained. Subsequently, the tables containing positive and negative enriched delta radiomic pathway pairs were aggregated, and rows and columns without significantly enriched results were removed, followed by visualization with the “pheatmap” R package.

### Correlation analysis

Spearman’s rank correlation coefficient rho was calculated between selected delta radiomic features and the log2-transformed expression intensity of every protein using base R packages, retaining only proteins with p<0.05 and ⍴≥0.6 for further analysis. Pearson’s correlation coefficient r was calculated between selected delta radiomic features and the fraction of α-SMA positive cells using base R packages.

### Cell Type Signature Enrichment Analysis

To infer relative cell type frequency changes between two groups from proteomics data, we applied signature enrichment analysis as described in ^30,31^, utilizing their published dataset. Cell type signatures were defined as sets of genes with cell-type specific gene expression of log2 fold change>0.3 and adjusted p<0.05 (**Supplementary Table 5**). For each cell type, we then tested for the enrichment in a ranked list of DE proteins (log2 fold changes) or correlation coefficients (weighted by -log10 p-value) using the Kolmogorov-Smirnow test. Positive and negative signed enrichment scores (-log10 p-values signed by effect size) reflect relative depletion and enrichment of the respective cell type, respectively. To visualize and interpret cell type signatures enriched in the the sets of proteins highly-correlation (Spearman’s |⍴|≥0.6, p<0.05) with delta radiomic features, the signed enrichment scores of each variable were transformed into a matrix with delta radiomic features as columns and cell types as rows. Enriched results with enrichment score<2 were dropped (set to zero), followed by removal of rows and columns without enriched entries, and visualization with the “pheatmap” R package.

### Association analysis with clinical parameters

Association analyses were performed to investigate associations of patient delta radiomics-derived (k-means) clusters with clinical parameters. Mann-Whitney U test was used for comparison of numerical variables, and Fisher’s exact test was used to compare categorical variables.

### Statistical analyses

All statistical analyses were performed in R (v.4.3.1.) environment. For all analyses, a p<0.05 was considered statistically significant unless stated otherwise. The following R packages were used: “readxl”, “clusterSim”, “dplyr”, “tidyverse”, “xlsx”, “viridis”, “hopkins”, “seriation”, “factoextra”, “RColorBrewer”, “pheatmap”, “fpc”, “caret”, “rstatix”, “pROC”, “ggpubr”, “ggpmisc”, “ggrepel”, “psych”, “heatmaply”, “UniProt.ws”, “AnnotationDbi”, “org.Mm.eg.db”, “limma”, “clusterProfiler”, “ReactomePA”, “parallel”, “doParallel”, “msigdbr”, “scales”, “DOSE”.

### Data Visualization

Figures were created in Adobe Illustrator (v.28.2) and partially contain graphics or illustrations from Adobe Stock and BioRender.com accessed through the academic licenses of the University of Bern.

## Abbreviations

α-SMA: Alpha-smooth muscle actin
AT1: alveolar type I
AT2: alveolar type II
BP: Biological process (ontology)
CC: Cellular compartment (ontology)
CT: Computed tomography
CTD: Connective tissue disease
DAPI: 4’,6-diamidino-2-phenylindole
DE: Differentially expressed
DLCO: Diffusing capacity of the lung for carbon monoxide
DPBS: Dulbecco’s phosphate buffered saline
ECM: Extracellular matrix
FDR: False discovery rate
FEV: Forced expiratory volume in the first second
FVC: Forced vital capacity
GO: Gene Ontology
HP: Hypersensitivity pneumonitis
HRCT: High-resolution computed tomography
HU: Hounsfield unit
i.t.: Intratracheal
IBSI: Imaging biomarkers standardization initiative
ICC: Intraclass correlation coefficient
ILD: Interstitial lung disease
IPF: Idiopathic pulmonary fibrosis
IQR: Interquartile range
KAEA: Kinase activity enrichment analysis
MF: Molecular function (ontology)
mPAP: Mean pulmonary arterial pressure
MRI: Magnetic resonance imaging
p.o.: *per os* (orally)
P(A)H: Pulmonary (arterial) hypertension
PDPN: Podoplanin
PF-ILD: Progressive fibrosing interstitial lung disease
PFT: Pulmonary function test
proSP-C: Prosurfactant Protein C
q.d.: *quaque die* (once a day)
QLF: Quantitative lung fibrosis
RTK: Receptor tyrosine kinase
scRNA-seq: Single-cell RNA sequencing
SSc: Systemic sclerosis

## Supplementary Tables

**Supplementary Table 1.** Summary of HRCT scan acquisition parameters.

| CT Parameter | Description |
| --- | --- |
| Manufacturer(s)* | Siemens, Siemens Healthineers, Philipps, Toshiba |
| Acquisition mode | Inspiration (breath hold) |
| Position | head first-supine (HFS), feet first-supine (FFS) |
| Slice thickness (mm) | 1 [range 0.5-1.5] |
| Reconstruction kernels | I70f, I80s, Br56f, Br56u, Br59f, LUNG, FC55 (sharp), YB |
| Tube voltage (kVp) | 120 (range 80-130) |

\*HRCT scanner models included SOMATOM Definition Flash (Siemens), SOMATOM Definition Edge (Siemens), SOMATOM Scope (Siemens) NAEOTOM Alpha (Siemens Healthineers), Aquilion (Toshiba), Brilliance 64 (Philips)

**Supplementary Table 2.**
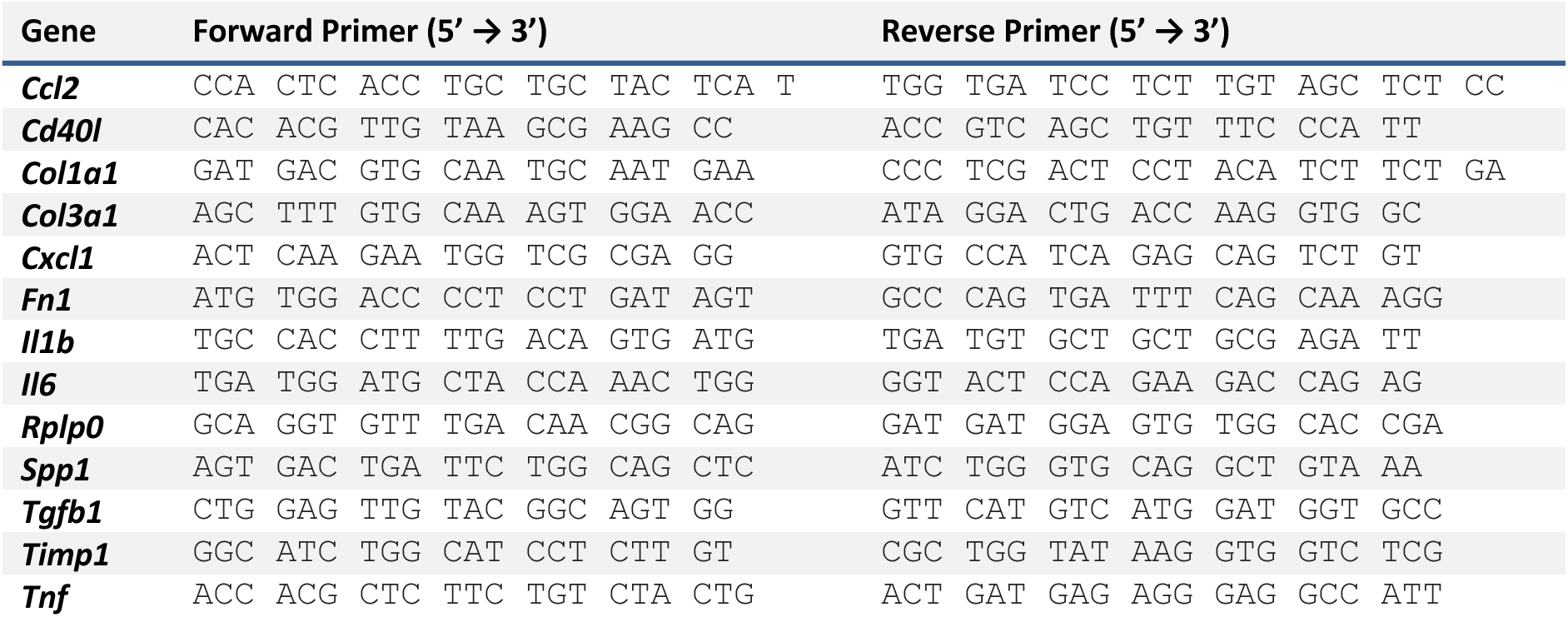
Mouse primer sequences used for quantitative PCR reactions.

**Supplementary Table 3.** Primary and secondary antibodies used for immunofluorescence staining.

| Description | Target | Reactivity | Host | Company | Label | Catalog | Dilution |
| --- | --- | --- | --- | --- | --- | --- | --- |
| Primary Ab | α-SMA | ms (hu, rt) | ms | Sigma-Aldrich | unconjugated | A5228 | 1:1500 |
| Primary Ab | proSP-C | ms (hu, rt) | rb | Merck-Millipore | unconjugated | AB3786 | 1:200 |
| Primary Ab | Podoplanin | ms | gt | R&D Systems | unconjugated | AF3244 | 1:200 |
| Secondary Ab | IgG | ms | dk | Invitrogen | Alexa488 | A21202 | 1:250 |
| Secondary Ab | IgG | rb | dk | Invitrogen | Alexa568 | A10042 | 1:250 |
| Secondary Ab | IgG | gt | dk | Invitrogen | Alexa647 | A21447 | 1:250 |

Abbreviations: ms, mouse; hu, human; rt, rat; rb, rabbit; gt, goat. Abbreviation: Ab, antibody.

**Supplementary Table 4.** Variable importance selected delta radiomic features used to classify treatment response.

| # | Feature Name | Feature Class | Feature Subclass | Filter | Score |
| --- | --- | --- | --- | --- | --- |
| 1 | *GLSZM_smallZone_lowGL_emp* | Texture | Texture | none | 0.92 |
| 2 | *LHL_GLSZM_GLvar* | Wavelet | Texture-derived | LHL | 0.96 |
| 3 | *center_mass_shift* | Shape | Shape | none | 1.00 |
| 4 | *LLL_hist_skewness* | Wavelet | Intensity-derived | LLL | 1.00 |
| 5 | *HLH_mGLCM_clust_prominence* | Wavelet | Texture-derived | HLH | 0.92 |
| 6 | *LHL_GLRLM_entropy* | Wavelet | Texture-derived | LHL | 0.92 |
| 7 | *HHL_GLRLM_runPercentage* | Wavelet | Texture-derived | HHL | 0.92 |
| 8 | *LHH_GLRLM_shortRunEmp* | Wavelet | Texture-derived | LHH | 1.00 |
| 9 | *LLH_mGLCM_IMC1* | Wavelet | Texture-derived | LLH | 0.96 |
| 10 | *HLL_hist_energy* | Wavelet | Intensity-derived | HLL | 0.96 |
| 11 | *LHH_mGLRLM_runPercentage* | Wavelet | Texture-derived | LHH | 0.92 |
| 12 | *LLL_GLSZM_largeZone_highGL_emp* | Wavelet | Texture-derived | LLL | 0.92 |
| 13 | *LHL_mGLCM_IMC2* | Wavelet | Texture-derived | LHL | 1.00 |
| 14 | *LHL_mGLCM_IMC1* | Wavelet | Texture-derived | LHL | 1.00 |
| 15 | *HLH_GLRLM_entropy* | Wavelet | Texture-derived | HLH | 0.96 |
| 16 | *LLH_mGLCM_MCC* | Wavelet | Texture-derived | LLH | 0.92 |
| 17 | *LLH_mGLCM_clust_prominence* | Wavelet | Texture-derived | LLH | 1.00 |
| 18 | *HLL_mGLCM_correlation* | Wavelet | Texture-derived | HLL | 0.96 |
| 19 | *HLH_hist_energy* | Wavelet | Intensity-derived | HLH | 0.92 |
| 20 | *LHL_mGLCM_MCC* | Wavelet | Texture-derived | LHL | 1.00 |
| 21 | *LLL_hist_coeffOfVar* | Wavelet | Intensity-derived | LLL | 1.00 |
| 22 | *HHL_mGLCM_IMC1* | Wavelet | Texture-derived | HHL | 0.92 |
| 23 | *LLH_GLSZM_ZSnonuniformity* | Wavelet | Texture-derived | LLH | 0.96 |
| 24 | *LLL_GLSZM_GLvar* | Wavelet | Texture-derived | LLL | 0.96 |
| 25 | *LHL_GLDZM_ZSnonuniformity_norm* | Wavelet | Texture-derived | LHL | 0.92 |
| 26 | *LHL_mGLCM_homogeneity_norm* | Wavelet | Texture-derived | LHL | 0.92 |
| 27 | *LLH_hist_energy* | Wavelet | Intensity-derived | LLH | 1.00 |
| 28 | *LLL_hist_kurtosis* | Wavelet | Intensity-derived | LLL | 0.92 |
| 29 | *LLH_GLSZM_GLnonuniformity_norm* | Wavelet | Texture-derived | LLH | 1.00 |
| 30 | *LLL_hist_percentile90* | Wavelet | Intensity-derived | LLL | 0.92 |
| 31 | *LHH_mGLCM_IMC2* | Wavelet | Texture-derived | LHH | 1.00 |
| 32 | *LHL_GLCM_MCC* | Wavelet | Texture-derived | LHL | 0.92 |
| 33 | *HHH_GLCM_IMC1* | Wavelet | Texture-derived | HHH | 1.00 |
| 34 | *HHH_GLRLM_LRvar* | Wavelet | Texture-derived | HHH | 1.00 |
| 35 | *GLSZM_smallZoneEmp* | Texture | Texture | none | 0.98 |
| 36 | *LHH_mGLCM_IMC1* | Wavelet | Texture-derived | LHH | 1.00 |
| 37 | *HHH_mGLRLM_entropy* | Wavelet | Texture-derived | HHH | 1.00 |
| 38 | *LHL_hist_energy* | Wavelet | Intensity-derived | LHL | 1.00 |
| 39 | *LHH_mGLRLM_GLnonuniformity_norm* | Wavelet | Texture-derived | LHH | 1.00 |
| 40 | *HHL_mGLRLM_entropy* | Wavelet | Texture-derived | HHL | 0.96 |
| 41 | *LHL_NGTDm_contrast* | Wavelet | Texture-derived | LHL | 0.92 |
| 42 | *LLH_GLCM_MCC* | Wavelet | Texture-derived | LLH | 0.92 |
| 43 | *LHL_GLSZM_ZSnonuniformity* | Wavelet | Texture-derived | LHL | 0.96 |
| 44 | *HHH_NGTDm_busyness* | Wavelet | Texture-derived | HHH | 1.00 |
| 45 | *HHH_mGLCM_MCC* | Wavelet | Texture-derived | HHH | 1.00 |
| 46 | *HLH_GLSZM_ZSnonuniformity* | Wavelet | Texture-derived | HLH | 1.00 |
| 47 | *LHH_GLCM_IMC1* | Wavelet | Texture-derived | LHH | 0.94 |
| 48 | *LHL_GLDZM_ZSnonuniformity* | Wavelet | Texture-derived | LHL | 1.00 |
| 49 | *HHH_GLRLM_longRun_lowGL_emp* | Wavelet | Texture-derived | HHH | 0.96 |
| 50 | *HHH_NGTDm_strength* | Wavelet | Texture-derived | HHH | 1.00 |
| 51 | *HHH_NGTDm_complexity* | Wavelet | Texture-derived | HHH | 1.00 |
| 52 | *LLH_GLSZM_entropy* | Wavelet | Texture-derived | LLH | 0.92 |
| 53 | *HHH_GLDZM_largeDistance_highGL_emp* | Wavelet | Texture-derived | HHH | 0.92 |
| 54 | *LHH_GLCM_MCC* | Wavelet | Texture-derived | LHH | 0.96 |

Abbreviations: GLCM = Gray Level Co-occurrence Matrix, NGTDM = Neighborhood Gray Tone Difference Matrix, GLRLM = Gray Level Run Length Matrix, GLDZM = Gray Level Distance Matrix and NGLDM = Neighboring Gray Level Dependence Matrix.

**Supplementary Table 5.** Cell type signature marker table used for cell type signature enrichment in bulk proteomics data. Data was adapted from ^30^. Only markers exhibiting |log_2_FC|>0.3 and FDR-adjusted p<0.05 were considered. Each entry is described by gene symbol, p-value, average log_2_FC, percentage of cells where the gene is detected in the cluster for the assigned cluster (pct.1) and in the other cluster(s) (pct.2), adjusted p-value, cell type annotation, and meta cell type annotation. Table provided as XLSX file. Data available in Supplementary XLSX File.

**Supplementary Table 6.** Differentially expressed proteins in nintedanib-treated mice in cluster 1 compared to vehicle-treated mice. The list includes all identified 7006 proteins. Only proteins with |log_2_FC|>0.3 and p<0.05 were considered for analysis. Each entry is described by log_2_FC, confidence intervals, average expression, t-value, p-value, B-value, Entrez ID, and gene symbol. Table provided as XLSX file. Data available in Supplementary XLSX File.

**Supplementary Table 7.** Differentially expressed proteins in nintedanib-treated mice in cluster 2 compared to vehicle-treated mice. The list includes all identified 7006 proteins. Only proteins with |log_2_FC|>0.3 and p<0.05 were considered for analysis. Each entry is described by log_2_FC, confidence intervals, average expression, t-value, p-value, B-value, Entrez ID, and gene symbol. Table provided as XLSX file. Data available in Supplementary XLSX File.

**Supplementary Table 8.** Differentially expressed proteins in nintedanib-treated mice in cluster 1 compared to nintedanib-treated mice in cluster 2. The list includes all identified 7006 proteins. Only proteins with |log_2_FC|>0.3 and p<0.05 were considered for analysis. Each entry is described by log_2_FC, confidence intervals, average expression, t-value, p-value, B-value, Entrez ID, and gene symbol. Table provided as XLSX file. Data available in Supplementary XLSX File.

**Supplementary Table 9.** Results of Gene Ontology enrichment analysis of differentially expressed, downregulated proteins (log_2_FC<-0.3, p<0.05) of nintedanib-treated mice in cluster 1 compared to nintedanib-treated mice in cluster 2. Each entry is described by ontology (BP, CC, or MF), GO identifier, pathway description, GeneRatio, BgRatio, p-value, FDR-adjusted p-value, q-value, and gene symbols. Table provided as XLSX file. Data available in Supplementary XLSX File.

**Supplementary Table 10.** Results of Gene Ontology enrichment analysis of differentially expressed, upregulated proteins (log_2_FC>0.3, p<0.05) of nintedanib-treated mice in cluster 1 compared to nintedanib-treated mice in cluster 2. Each entry is described by ontology (BP, CC, or MF), GO identifier, pathway description, GeneRatio, BgRatio, p-value, FDR-adjusted p-value, q-value, and gene symbols. Table provided as XLSX file. Data available in Supplementary XLSX File.

**Supplementary Table 11.** Result of Reactome enrichment analysis of the 54 radioproteomic association modules separated by annotation of proteins positively (Spearman’s ⍴≥0.6, p<0.05) and proteins negatively (Spearman’s ⍴≤-0.6, p<0.05) correlating with the preclinical response-defining delta radiomic feature. Each entry is described by delta radiomic feature name, protein subset entered into enrichment analysis, Reactome identifier, pathway description, GeneRatio, BgRatio, p-value, FDR-adjusted p-value, q-value, gene symbols, and count. Data available in Supplementary XLSX File.

**Supplementary Table 12.** Result of Gene Ontology - Biological Process (GO:BP) enrichment analysis of the 54 radioproteomic association modules separated by annotation of proteins positively (Spearman’s ⍴≥0.6, p<0.05) and proteins negatively (Spearman’s ⍴≤-0.6, p<0.05) correlating with the preclinical response-defining delta radiomic feature. Each entry is described by delta radiomic feature name, protein subset entered into enrichment analysis, GO identifier, pathway description, GeneRatio, BgRatio, p-value, FDR-adjusted p-value, q-value, gene symbols, and count. Data available in Supplementary XLSX File.

**Supplementary Table 13.** Results of cell type enrichment analysis of the 54 radioproteomic association modules. Proteins correlating (Spearman’s |⍴|≥0.6, p<0.05) with delta radiomic features were ranked by log10 p-value and weighted by correlation coefficient prior to entering into deconvolution analysis. Each entry is described by delta radiomic feature name, p-value, fold change difference, signed log10 enrichment p-value, cell type, and enrichment trend. Data available in Supplementary XLSX File.

**Supplementary Table 14.**
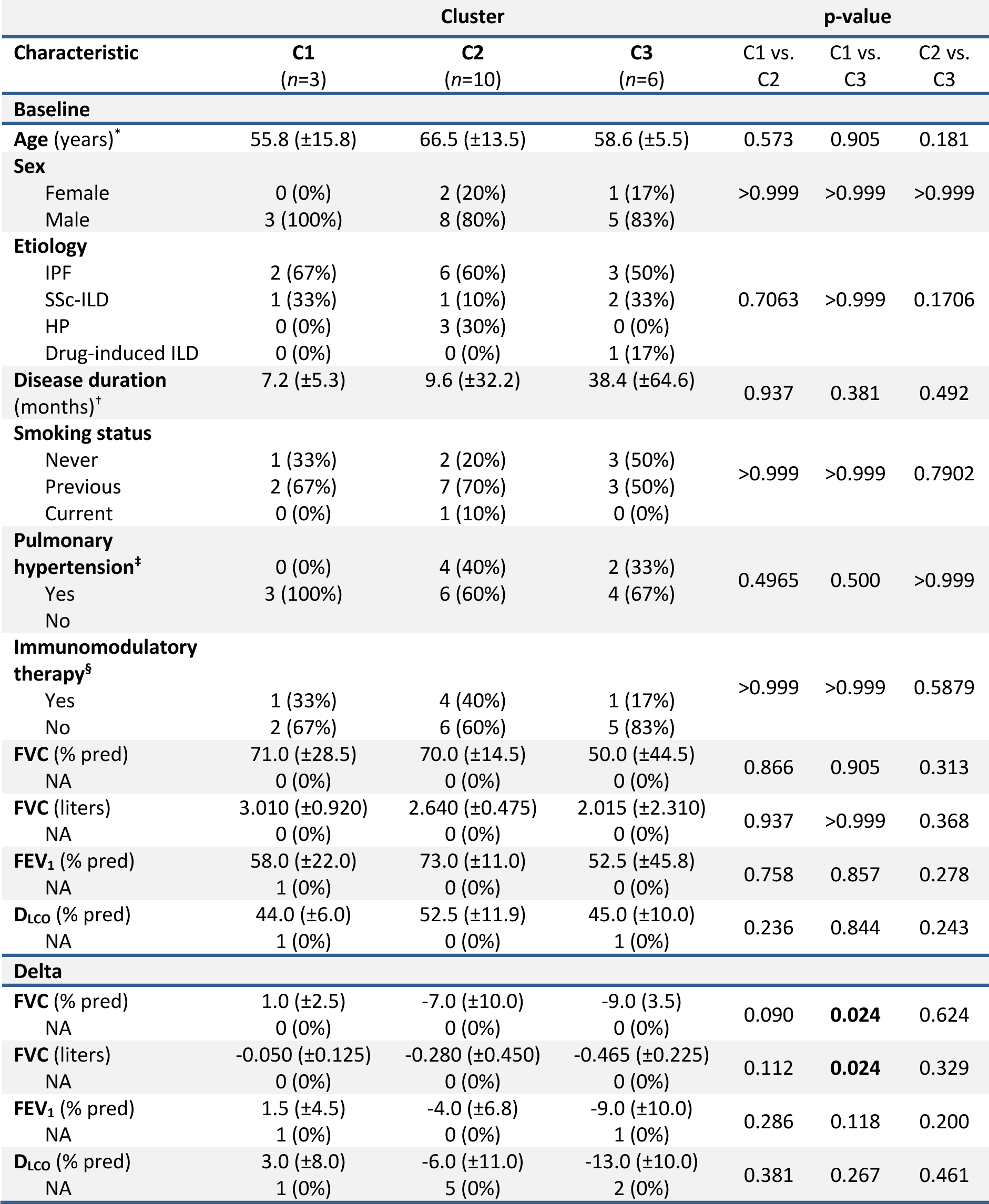
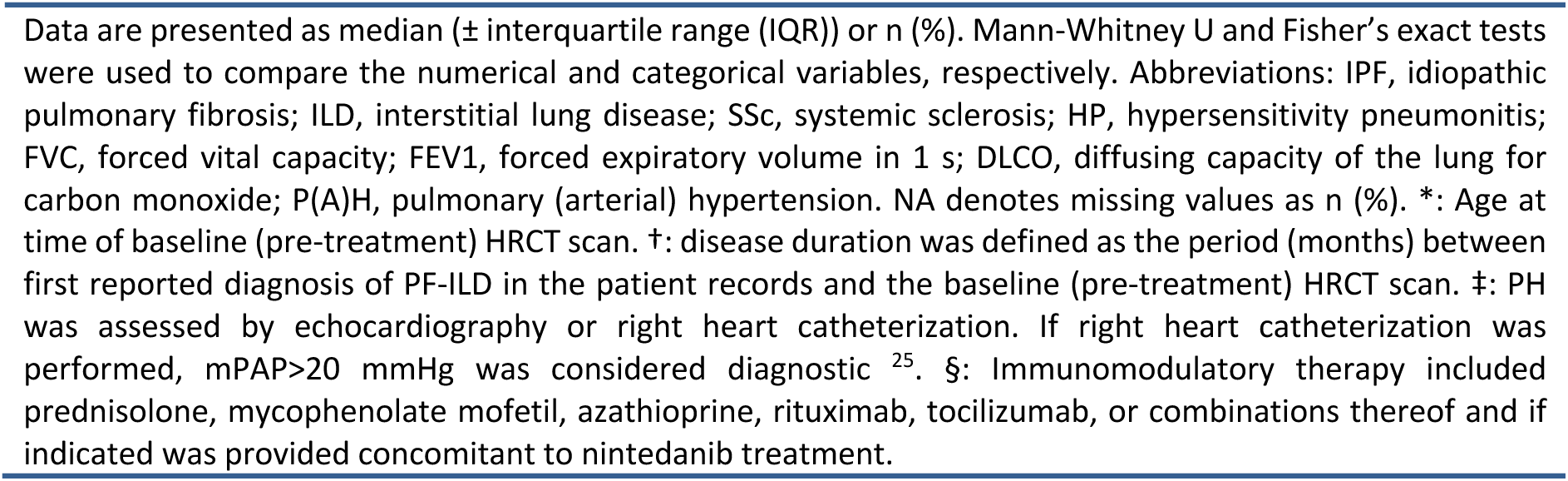
Associations of patients’ groups resulting from unsupervised clustering on preclinical treatment response-defining delta radiomic features (*n*=54) with clinical parameters.

|  |  |  |  |  |  |  |
| --- | --- | --- | --- | --- | --- | --- |
|  | **Cluster** |  |  | **p-value** |  |  |
| **Characteristic** | **C1**<br>($n=3$) | **C2**<br>($n=10$) | **C3**<br>($n=6$) | C1 vs.<br>C2 | C1 vs.<br>C3 | C2 vs.<br>C3 |
| **Baseline** |  |  |  |  |  |  |
| **Age (years)\*** | 55.8 ($\pm 15.8$) | 66.5 ($\pm 13.5$) | 58.6 ($\pm 5.5$) | 0.573 | 0.905 | 0.181 |
| **Sex** |  |  |  |  |  |  |
| Female | 0 (0%) | 2 (20%) | 1 (17%) | >0.999 | >0.999 | >0.999 |
| Male | 3 (100%) | 8 (80%) | 5 (83%) |  |  |  |
| **Etiology** |  |  |  |  |  |  |
| IPF | 2 (67%) | 6 (60%) | 3 (50%) |  |  |  |
| SSc-ILD | 1 (33%) | 1 (10%) | 2 (33%) | 0.7063 | >0.999 | 0.1706 |
| HP | 0 (0%) | 3 (30%) | 0 (0%) |  |  |  |
| Drug-induced ILD | 0 (0%) | 0 (0%) | 1 (17%) |  |  |  |
| **Disease duration (months)<sup>†</sup>** | 7.2 ($\pm 5.3$) | 9.6 ($\pm 32.2$) | 38.4 ($\pm 64.6$) | 0.937 | 0.381 | 0.492 |
| **Smoking status** |  |  |  |  |  |  |
| Never | 1 (33%) | 2 (20%) | 3 (50%) | >0.999 | >0.999 | 0.7902 |
| Previous | 2 (67%) | 7 (70%) | 3 (50%) |  |  |  |
| Current | 0 (0%) | 1 (10%) | 0 (0%) |  |  |  |
| **Pulmonary hypertension<sup>‡</sup>** |  |  |  |  |  |  |
| Yes | 0 (0%) | 4 (40%) | 2 (33%) | 0.4965 | 0.500 | >0.999 |
| No | 3 (100%) | 6 (60%) | 4 (67%) |  |  |  |
| **Immunomodulatory therapy<sup>§</sup>** |  |  |  |  |  |  |
| Yes | 1 (33%) | 4 (40%) | 1 (17%) | >0.999 | >0.999 | 0.5879 |
| No | 2 (67%) | 6 (60%) | 5 (83%) |  |  |  |
| **FVC (% pred)** | 71.0 ($\pm 28.5$) | 70.0 ($\pm 14.5$) | 50.0 ($\pm 44.5$) | 0.866 | 0.905 | 0.313 |
| NA | 0 (0%) | 0 (0%) | 0 (0%) |  |  |  |
| **FVC (liters)** | 3.010 ($\pm 0.920$) | 2.640 ($\pm 0.475$) | 2.015 ($\pm 2.310$) | 0.937 | >0.999 | 0.368 |
| NA | 0 (0%) | 0 (0%) | 0 (0%) |  |  |  |
| **FEV<sub>1</sub> (% pred)** | 58.0 ($\pm 22.0$) | 73.0 ($\pm 11.0$) | 52.5 ($\pm 45.8$) | 0.758 | 0.857 | 0.278 |
| NA | 1 (0%) | 0 (0%) | 0 (0%) |  |  |  |
| **D<sub>Lco</sub> (% pred)** | 44.0 ($\pm 6.0$) | 52.5 ($\pm 11.9$) | 45.0 ($\pm 10.0$) | 0.236 | 0.844 | 0.243 |
| NA | 1 (0%) | 0 (0%) | 1 (0%) |  |  |  |
| **Delta** |  |  |  |  |  |  |
| **FVC (% pred)** | 1.0 ($\pm 2.5$) | -7.0 ($\pm 10.0$) | -9.0 (3.5) | 0.090 | **0.024** | 0.624 |
| NA | 0 (0%) | 0 (0%) | 0 (0%) |  |  |  |
| **FVC (liters)** | -0.050 ($\pm 0.125$) | -0.280 ($\pm 0.450$) | -0.465 ($\pm 0.225$) | 0.112 | **0.024** | 0.329 |
| NA | 0 (0%) | 0 (0%) | 0 (0%) |  |  |  |
| **FEV<sub>1</sub> (% pred)** | 1.5 ($\pm 4.5$) | -4.0 ($\pm 6.8$) | -9.0 ($\pm 10.0$) | 0.286 | 0.118 | 0.200 |
| NA | 1 (0%) | 0 (0%) | 1 (0%) |  |  |  |
| **D<sub>Lco</sub> (% pred)** | 3.0 ($\pm 8.0$) | -6.0 ($\pm 11.0$) | -13.0 ($\pm 10.0$) | 0.381 | 0.267 | 0.461 |
| NA | 1 (0%) | 5 (0%) | 2 (0%) |  |  |  |

Data are presented as median ( ± interquartile range (IQR)) or n (%). Mann-Whitney U and Fisher's exact tests were used to compare the numerical and categorical variables, respectively. Abbreviations: IPF, idiopathic pulmonary fibrosis; ILD, interstitial lung disease; SSc, systemic sclerosis; HP, hypersensitivity pneumonitis; FVC, forced vital capacity; FEV<sub>1</sub>, forced expiratory volume in 1 s; DLCO, diffusing capacity of the lung for carbon monoxide; P(A)H, pulmonary (arterial) hypertension. NA denotes missing values as n (%). \*: Age at time of baseline (pre-treatment) HRCT scan. †: disease duration was defined as the period (months) between first reported diagnosis of PF-ILD in the patient records and the baseline (pre-treatment) HRCT scan. ‡: PH was assessed by echocardiography or right heart catheterization. If right heart catheterization was performed, mPAP>20 mmHg was considered diagnostic <sup>25</sup>. §: Immunomodulatory therapy included prednisolone, mycophenolate mofetil, azathioprine, rituximab, tocilizumab, or combinations thereof and if indicated was provided concomitant to nintedanib treatment.

**Supplementary Table 15.**
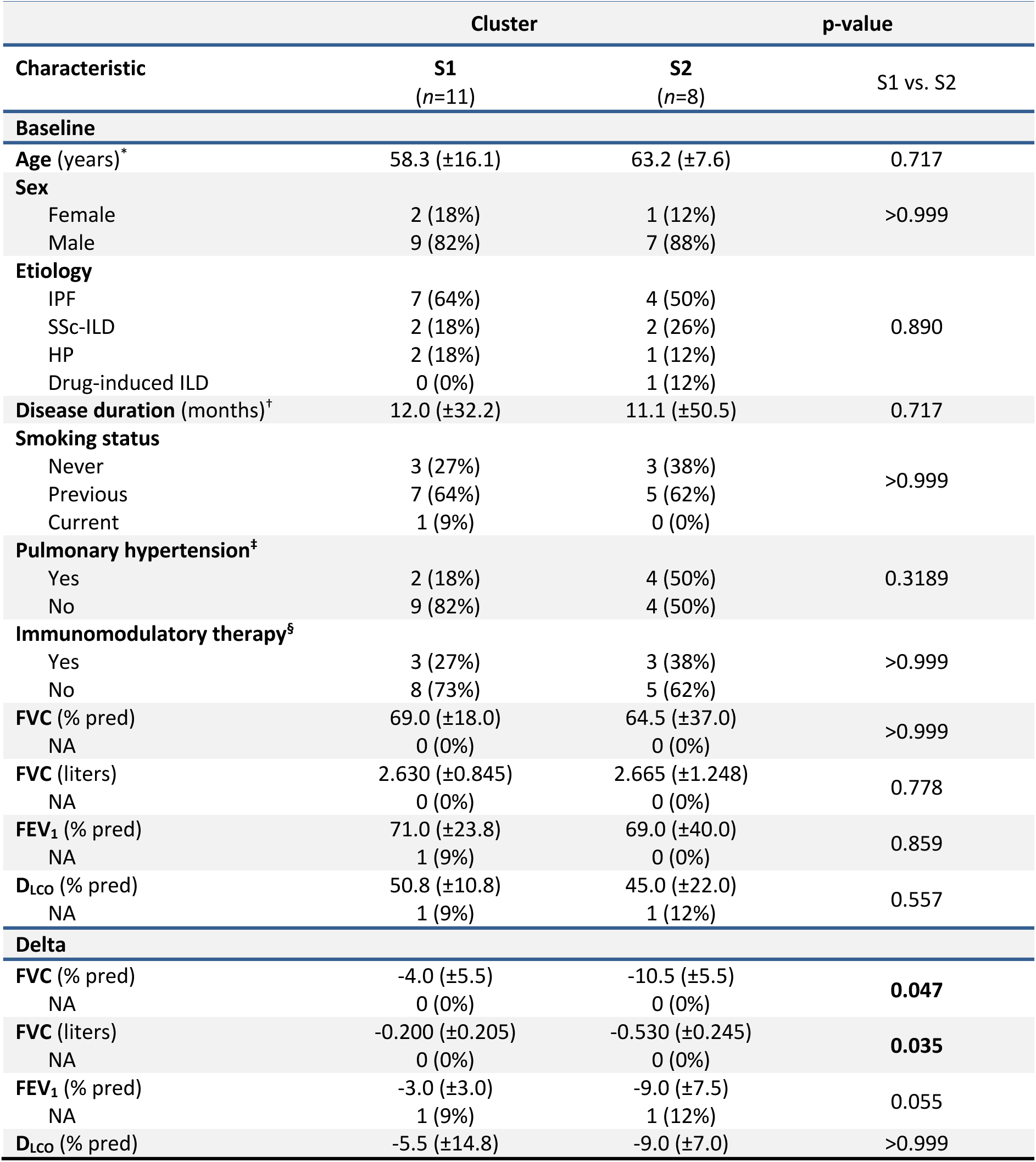
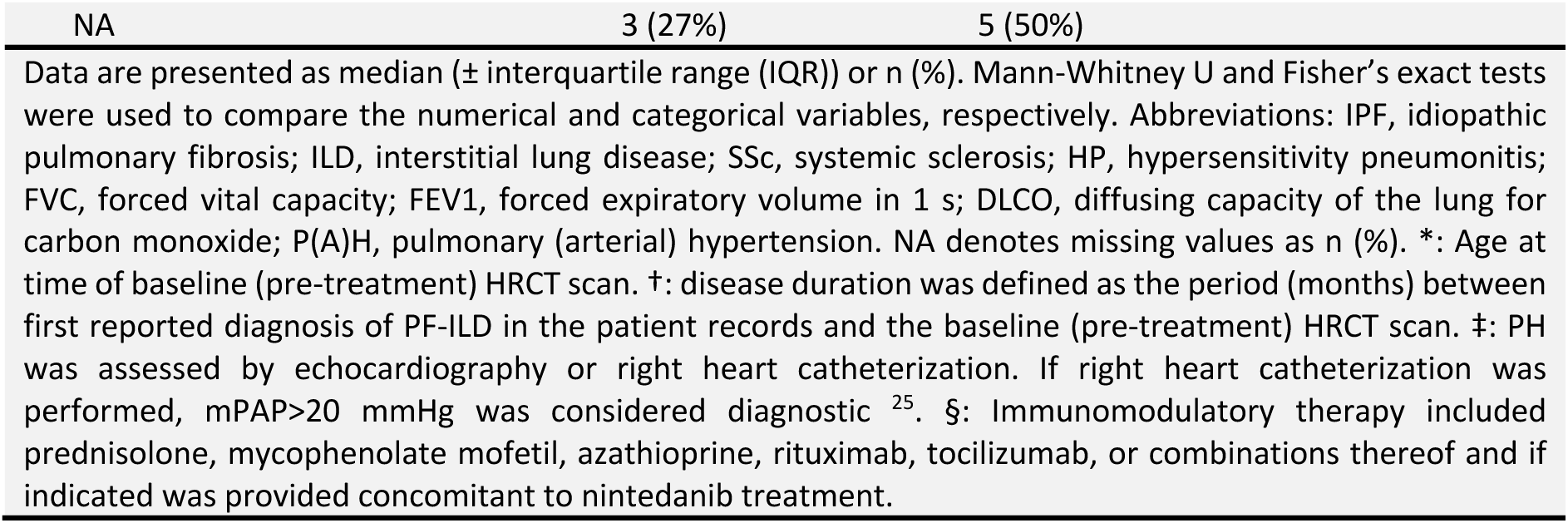
Associations of patients’ groups resulting from unsupervised clustering on ECM remodeling-associated delta radiomic features (*n*=8) with clinical parameters.

|  | Cluster |  | p-value |
| --- | --- | --- | --- |
| Characteristic | S1<br>(n=11) | S2<br>(n=8) | S1 vs. S2 |
| Baseline |  |  |  |
| Age (years)* | 58.3 (±16.1) | 63.2 (±7.6) | 0.717 |
| Sex |  |  |  |
| Female | 2 (18%) | 1 (12%) | >0.999 |
| Male | 9 (82%) | 7 (88%) |  |
| Etiology |  |  |  |
| IPF | 7 (64%) | 4 (50%) | 0.890 |
| SSc-ILD | 2 (18%) | 2 (26%) |  |
| HP | 2 (18%) | 1 (12%) |  |
| Drug-induced ILD | 0 (0%) | 1 (12%) |  |
| Disease duration (months)† | 12.0 (±32.2) | 11.1 (±50.5) | 0.717 |
| Smoking status |  |  |  |
| Never | 3 (27%) | 3 (38%) | >0.999 |
| Previous | 7 (64%) | 5 (62%) |  |
| Current | 1 (9%) | 0 (0%) |  |
| Pulmonary hypertension‡ |  |  |  |
| Yes | 2 (18%) | 4 (50%) | 0.3189 |
| No | 9 (82%) | 4 (50%) |  |
| Immunomodulatory therapy§ |  |  |  |
| Yes | 3 (27%) | 3 (38%) | >0.999 |
| No | 8 (73%) | 5 (62%) |  |
| FVC (% pred) | 69.0 (±18.0) | 64.5 (±37.0) | >0.999 |
| NA | 0 (0%) | 0 (0%) |  |
| FVC (liters) | 2.630 (±0.845) | 2.665 (±1.248) | 0.778 |
| NA | 0 (0%) | 0 (0%) |  |
| FEV1 (% pred) | 71.0 (±23.8) | 69.0 (±40.0) | 0.859 |
| NA | 1 (9%) | 0 (0%) |  |
| Dlco (% pred) | 50.8 (±10.8) | 45.0 (±22.0) | 0.557 |
| NA | 1 (9%) | 1 (12%) |  |
| Delta |  |  |  |
| FVC (% pred) | -4.0 (±5.5) | -10.5 (±5.5) | 0.047 |
| NA | 0 (0%) | 0 (0%) |  |
| FVC (liters) | -0.200 (±0.205) | -0.530 (±0.245) | 0.035 |
| NA | 0 (0%) | 0 (0%) |  |
| FEV1 (% pred) | -3.0 (±3.0) | -9.0 (±7.5) | 0.055 |
| NA | 1 (9%) | 1 (12%) |  |
| Dlco (% pred) | -5.5 (±14.8) | -9.0 (±7.0) | >0.999 |

| NA | 3 (27%) | 5 (50%) |
| --- | --- | --- |
| Data are presented as median (± interquartile range (IQR)) or n (%). Mann-Whitney U and Fisher's exact tests were used to compare the numerical and categorical variables, respectively. Abbreviations: IPF, idiopathic pulmonary fibrosis; ILD, interstitial lung disease; SSc, systemic sclerosis; HP, hypersensitivity pneumonitis; FVC, forced vital capacity; FEV1, forced expiratory volume in 1 s; DLCO, diffusing capacity of the lung for carbon monoxide; P(A)H, pulmonary (arterial) hypertension. NA denotes missing values as n (%). *: Age at time of baseline (pre-treatment) HRCT scan. †: disease duration was defined as the period (months) between first reported diagnosis of PF-ILD in the patient records and the baseline (pre-treatment) HRCT scan. ‡: PH was assessed by echocardiography or right heart catheterization. If right heart catheterization was performed, mPAP>20 mmHg was considered diagnostic <sup>25</sup>. §: Immunomodulatory therapy included prednisolone, mycophenolate mofetil, azathioprine, rituximab, tocilizumab, or combinations thereof and if indicated was provided concomitant to nintedanib treatment. |  |  |

